# Epigenetic deprogramming driven by disruption of CIZ1-RNA nuclear assemblies in early-stage breast cancers

**DOI:** 10.1101/2024.08.22.609161

**Authors:** Gabrielle L. Turvey, Ernesto López de Alba, Emma Stewart, Heather Cook, Ahmad Alalti, Richard T. Gawne, Justin F-X Ainscough, Andrew S. Mason, Dawn Coverley

**Affiliations:** Mammalian cell cycle research group, Department of Biology, University of York, YO10 5DD, UK; York Biomedical Research Institute, University of York, YO10 5DD, UK; Jack Birch Unit for Molecular Carcinogenesis, Department of Biology, University of York, YO10 5DD, UK; Topology and DNA breaks Group, Spanish National Cancer Centre (CNIO), Madrid 28029, Spain; Nuffield Department of Medicine, University of Oxford OX3 7FZ, UK

**Keywords:** CIZ1, Cancer, X chromosome, RNA-protein complexes, Chromatin

## Abstract

CIZ1 is part of the large RNA-dependent supramolecular assemblies that form around the inactive X-chromosome (Xi) in female cells, and smaller assemblies throughout the nucleus in males and females. Here we show that CIZ1 C-terminal anchor domain (AD) is elevated in primary human breast tumour transcriptomes, even at stage I. Elevation of AD correlates with deprotection of chromatin, and up-regulation of lncRNA-containing gene clusters in approximately 10Mb regions that are enriched in cancer-associated genes. We modelled the effect of ectopic AD on endogenous CIZ1-Xi assemblies and observed dominant-negative interference with their re-formation after mitosis, leading to abnormal Xi assemblies similar to those in breast cancer cells, and depletion of histone modifications H2AK119ub1 and H3K27me3. Within days consistent alterations in gene expression were evident across the genome, showing that disruption of CIZ1-RNA assemblies has a destabilizing effect, likely by unscheduled exposure of underlying chromatin to modifying enzymes. Together the data argue for a dominant, potent and rapid effect of CIZ1 AD, that can deprogram established patterns of gene expression and which may predispose incipient tumours to epigenetic instability.

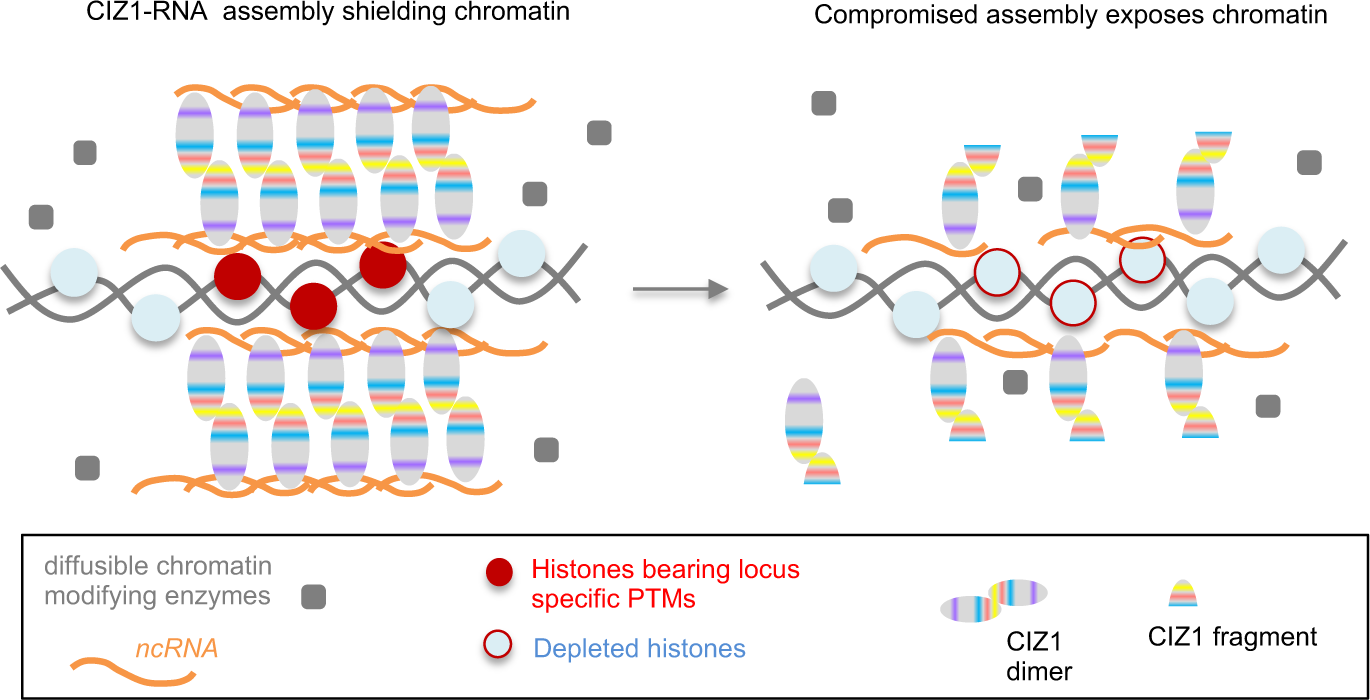

## Introduction

Selection and packaging of chromatin into transcriptionally repressed states underlies cell specialisation and development. Weakened repression of heterochromatin can result in pro-oncogenic changes, and has the potential to give rise to all the classic hallmarks of cancer, even in the absence of genetic change^1–3^. The inactive X chromosome (Xi) is the most intensely studied model of chromatin repression, revealing how the *cis*-acting lncRNA *Xist*^4, 5^ directs the formation of large RNA-dependent supramolecular assembly complexes (SMACs) populated by chromatin modifying enzymes^6^. Aggregation of SMAC proteins, mediated by their intrinsically disordered regions (IDRs), creates a functional nuclear compartment that partitions regulatory factors to establish local gene silencing early in development.

Cip1-interacting zinc finger protein 1 (CIZ1) is one of several proteins that populate Xi SMACs, recruited via its interaction with the repeat E element of *Xist*^7, 8^. Several observations set CIZ1 apart from other SMAC components. First, it is not required for *Xist* recruitment, Xi silencing or embryonic development, and the impact of its loss only becomes apparent in somatic cells in which repressed chromatin is already established, but must be faithfully maintained. A requirement for CIZ1 is apparent in differentiated fibroblasts from CIZ1 null mice, where local retention of *Xist* around Xi chromatin is compromised^7, 8^, repressive histone post-translational modifications (PTMs) are lost, and genome-wide changes in the expression of genes under the regulation of polycomb repressive complexes (PRC 1 and 2) are apparent^9^. Second, the stability of CIZ1 within Xi SMACs, even those that form during the initiation stages of X-inactivation, is unusually high. Compared to other SMAC components the residency time of CIZ1 is estimated to be 2-10 fold longer, similar to that of *Xist*^6^. Thus, it appears that CIZ1 exchanges less readily than other protein components, and might therefore contribute a stabilizing influence on *Xist* and Xi SMACs.

Some of the sequence determinants required for assembly of CIZ1 within Xi SMACs are known, including two alternatively-spliced, low-complexity prion-like domains (PLD1 and PLD2) that modulate interaction with *Xist,* and a second RNA interaction domain in the C-terminus ^10^. Neither RNA interaction is sufficient to support assembly of CIZ1 into Xi SMACs on its own, but together they drive both assembly, and the *de novo* enrichment of H2AK119ub1 and H3K27me3, added by PRC1 and 2 respectively, in underlying chromatin. These experiments directly link CIZ1 SMAC formation with modification of chromatin and implicate its bi-valent interaction with RNA^10^.

Disappearance of the Barr body (Xi), has been known for decades and is considered a hallmark of cancer^11^. Erosion of the Xi in breast tumours and cell lines was originally ascribed to genetic instability, though epigenetic instability is also apparent, evident as abnormal sub-nuclear organization, aberrant promoter DNA methylation, and perturbations of chromatin including H3K27me3^12^. Transcriptional reactivation of X-linked genes has been implicated in both breast and ovarian cancers^13^ though is likely to be indicative of wider, and possibly earlier, epigenetic erosion. In fact, widespread erosion of the DNA methylation landscape can give rise to the transcriptional changes common in tumors^14^, and for breast cancers in particular the progression from progenitor cell to pre-malignant lesion has been shown to involve changes in the DNA methylome that precede genetic instability^15, 16^. From data such as these, a model is emerging in which induction of breast cancer could occur primarily through epigenetic disruption.

Here we describe the aberrant expression of CIZ1 in human cancers, and model the effects of destabilising protein fragments on RNA-protein assemblies and underlying chromatin. The data lead to the conclusion that disease-associated dominant-negative CIZ1 fragments (DNFs) contribute to epigenetic instability by deprotecting loci that are normally buffered by surrounding SMACs, and that this plays an early role in tumour aetiology by promoting epigenetic instability.

## Results

### CIZ1 assemblies are disrupted in breast cancer cells

In primary epithelial cells derived from normal human female mammary tissue (HMECs) a single large CIZ1 assembly is visible in approximately 80% of cells in a cycling population (Fig.1A). This coincides with local enrichment of H2AK119ub1 identifying the assembly as at the Xi, as reported for human^7, 17, 18^ and murine cells^6–8^. Human and murine CIZ1 possess the same conserved domains encoded by the same exons in the same order (SFig.1A), and so far no differences in behaviour or function have been uncovered. CIZ1-Xi assemblies are dependent on multivalent interaction with RNAs including *Xist* (Fig.1B^10^), and normally observed with similar frequency regardless of whether epitopes in its N-terminal DNA replication domain (RD^19^), or C-terminal nuclear matrix anchor domain (AD^20^) are detected (Fig.1C).

**Figure 1.**
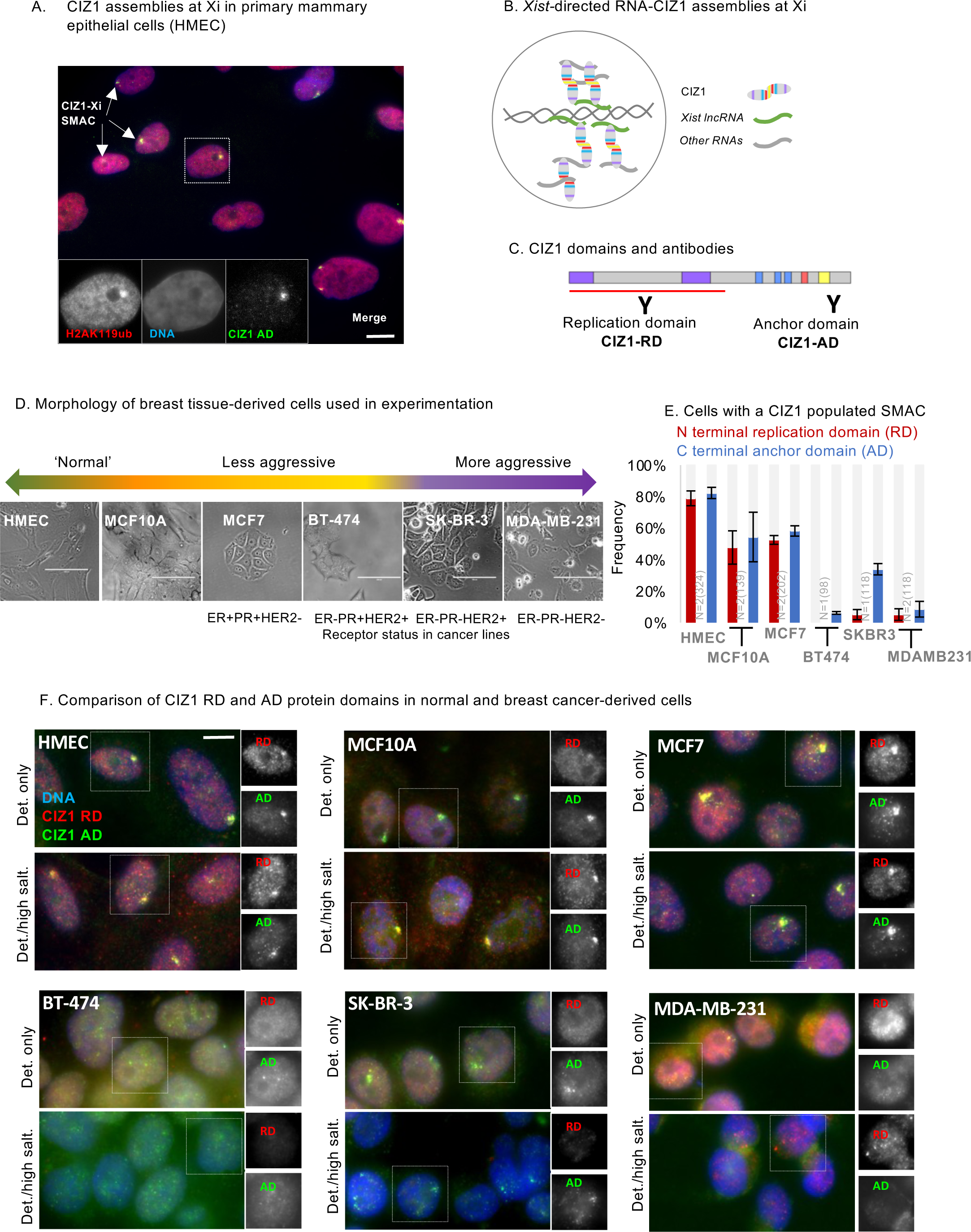
Corrupted CIZ1-Xi assemblies in breast cancer cells. A) Female primary human breast epithelial cells (HMECs) stained for CIZ1 via its C-terminal anchor domain (AD, green), and co-localization with H2AK119ub1 (red) as a marker of Xi chromatin. DNA is blue. Inset, example nucleus with CIZ1-AD and H2AK119ub1 shown individually in grey scale. Bar is 10μm. B) Model showing multivalent interaction between CIZ1 N- and C-terminal RNA interaction domains, and RNAs including *Xist* in the vicinity of the inactive X chromosome ^10^. C) Schematic of CIZ1 protein domains (see also SFig.1A) and location of epitopes used to report on CIZ1-AD (green, Ab87) or CIZ1 replication domain (RD, red, Ab1793). D) Bright field images of breast-derived cell types ordered based on phenotype, with corresponding hormone and growth factor receptor status. Bar is 100 microns. E) Frequency of cells with discrete nuclear CIZ1 assemblies, detected via CIZ1-AD or CIZ1-RD in cycling populations of the indicated breast-derived cell types. Error bars show SEM. A reduced frequency of CIZ1-Xi assemblies is observed in non-cancer breast cell line MCF-10A and cancer cell line MCF7, while in the more aggressive BT-474 and MDA-MB-231 cancer cells, large CIZ1 SMACs are rare for both RD and AD epitopes, and in SK-BR-3 populations only detectable via the AD epitope. In all four of the cancer lines the appearance of those assemblies that are detected is less compact and coherent. F) Example immunofluorescence images of CIZ1-RD (red) and CIZ1-AD (green) in HMEC and the indicated breast cancer cell lines, after pre-fixation wash with detergent-containing buffer (Det. only), or after high-salt extraction (Det./high salt). Right, nuclei in which RD and AD are shown individually in grey scale. Bar is 10μm. The RD and AD epitopes were differentially detected or extracted in some cases, indicating that they are not always part of the same polypeptide (for example compare nucleus-wide RD in SK-BR-3 cells, in detergent-treated cells to detergent/high-salt treated cells).

However, compared to normal breast tissue-derived cells or cell lines (Fig.1D), in breast cancer-derived cell lines CIZ1 Xi assemblies are either absent, less compact and coherent, or RD and AD epitopes are differentially susceptible to extraction from the nucleus (Fig.1E,F). This indicates that CIZ1 RD and AD are not always part of the same polypeptide, and are compromised in their ability to form stable assemblies around Xi chromatin. We conclude that CIZ1 protein expression, and CIZ1-Xi assemblies are commonly disrupted in breast cancer cell lines, consistent with the reported wider destabilization of the inactive X chromosome in breast cancer cells and tissues^12^.

Alignment of transcriptomes from four breast cancer-derived cell lines and a control cell line, to CIZ1’s translated exons (2-17) revealed over-representation of AD-encoding exons in the tumour-derived lines, compared to RD-encoding exons (SFig.1B, Supplemental Data set 1). We also noted a transition in transcript coverage within exon 10 (SFig.1C), which coincides with an internal transcription start sites (TSS) annotated in Ensembl^21^ from the FANTOM5 project^22^, and with enrichment of indicators of active chromatin in cancer cell lines but not normal HMECs (SFig.1D). Thus, archive data suggest that transcription can begin from an internal site in the CIZ1 gene.

### Elevation of CIZ1 AD-encoding transcript in early-stage primary breast tumours

To measure CIZ1 transcript expression in primary common solid tumours, we first used quantitative RT-PCR to compare the 5’ end to the 3’ end (which contribute coding sequence to RD and AD respectively) by detection of amplicons unaffected by alternative splicing ^23^ (Fig.2A). In cDNAs from 46 tissue samples (SFig.2A), correlation between two RD amplicons (in exons 5 and 7), or between two AD amplicons (in exons 14 and 16) was strong, however, RD and AD did not correlate with each other. This confirms that expression of RD and AD are commonly uncoupled at the transcript level, and shows that the differential can be sampled across exons 5-7 verses 14-16.

**Figure 2.**
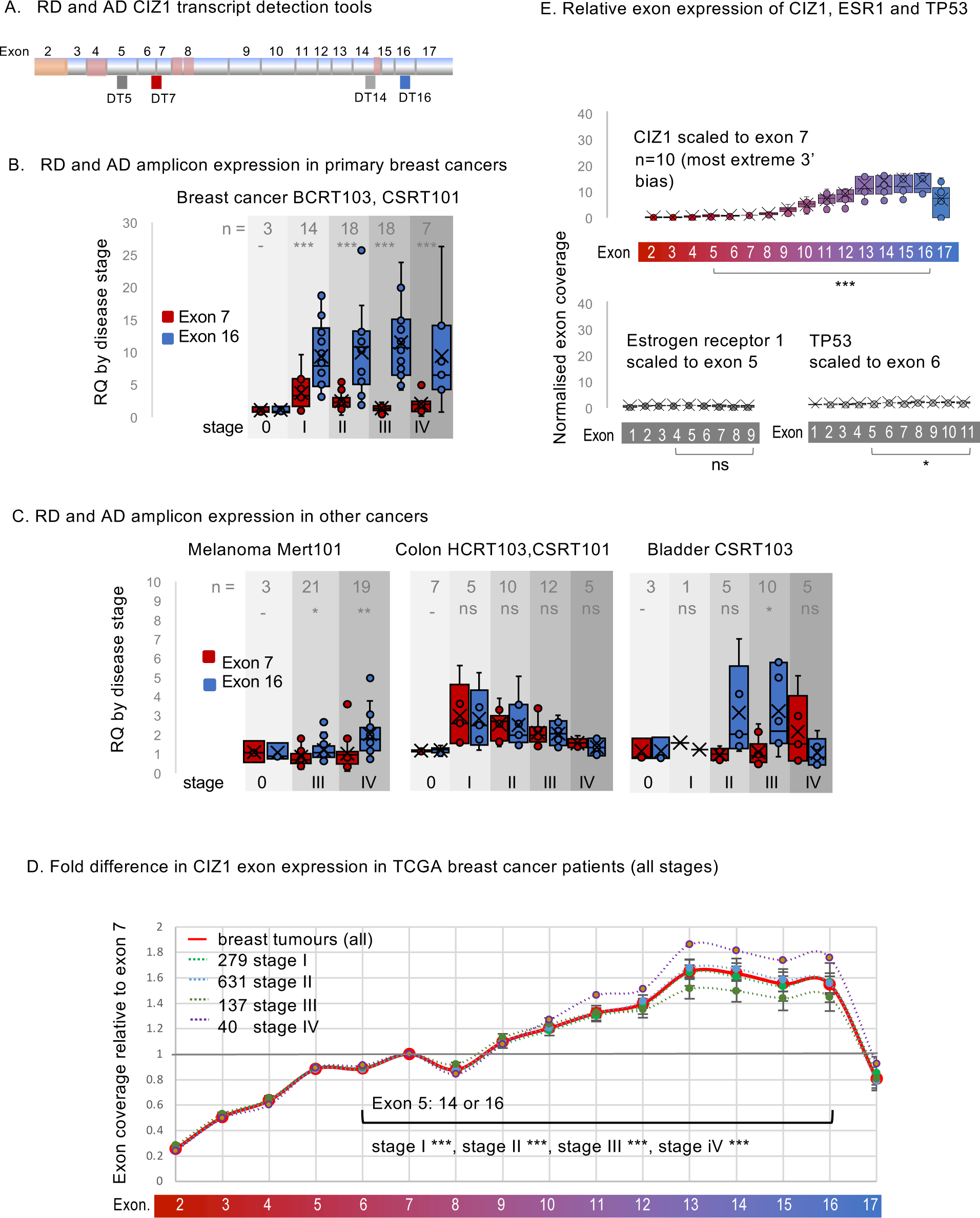
Elevated CIZ1 anchor domain expression in primary cancers. A) Exon structure of CIZ1 based on human reference sequence NM_012127.2 showing all 16 translated exons (2-17), and those subject to alternative splicing (pink)^10, 19, 55–58^. The location of amplicons detected by quantitative RT-PCR detection tools are shown (see also SFig.2A). Alternative untranslated exons 1’s are not shown. B) Relative quantification (RQ) of CIZ1 exon 7 (red) and CIZ1 exon 16 (blue) in primary human breast tissue-derived cDNAs in arrays BCRT103 and CSRT101 (n=60). Box and whisker plots show results aggregated by clinical stage (0-IV), calibrated to the average of the stage 0 samples for each amplicon where 0 represents histologically normal tissue. Significance indicators show comparisons between amplicons by t-test, where ns is not significant, *p<0.05, **p<0.01, ***p<0.001. Individual sample values are given in Supplemental Dataset 2. C) As in B for human tissue-derived cDNAs in arrays MERT101 (melanoma, n=43), HCRT103, CSRT101 (colon, n=39) and CSRT103 (bladder, n=24). D) CIZ1 exon expression in TCGA breast cancer samples, separated by clinical stage and normalised to individual exon 7 expression. At all stages, 5’ and 3’ expression is significantly different, with 3’ elevation from around exon 10. Comparison of transcript levels in exon 5, to 14 or 16 (arrows) is by Mann-Whitney U test. Error bars show SEM. E) Control analysis showing TPMs in a subset of 10 stage 2 TCGA breast cancer patients that exhibit the most marked 3’ end bias for CIZ1, mapped to CIZ1 exons, normalised to exon 7. Below, TPMs from the same patients for estrogen receptor alpha (ERα/ESR1) normalised to its exon 5, and TP53 normalised to its exon 6, showing relative exon coverage and lack of 3’ over-representation.

Domain disparity was striking and consistent in breast tumours across all stages (Fig.2B). It was also significant in bladder cancer at stage III and melanoma at stage III and IV (Fig.2C), and observed sporadically in other tumours of different aetiology (SFig.2B). In addition, in some colon, lung and thyroid tumours both RD and AD domains of CIZ1 were elevated, compared to histologically normal tissue (Fig.2C, SFig.2B, Supplemental Data set 2).

Focussing on breast cancer, we analysed *CIZ1* expression in 1095 transcriptomes submitted to The Cancer Genome Atlas (TCGA). While transcripts that map to the whole *CIZ1* gene revealed no *overall* difference in expression between tumours and normal tissue (SFig.2C), the same raw data when mapped to individual CIZ1 exons showed that AD (exon 14) is significantly over-represented compared to RD (exon 5) at all stages (Fig.2D, Supplemental Data set 1), and that AD elevation was notable from around exon 10. Similar elevation of C-terminal transcript was not evident in cancer-associated genes ESR1 and TP53 in a subset of the same transcriptomes (Fig. 2E). Together these data show that C-terminal CIZ1 exons are over-represented in the majority of breast cancers, that epitopes encoded by C-terminal exons are uncoupled from N-terminal exons, and pose the question of whether inappropriate AD protein is functionally relevant.

### *In vitro* modelling of the effect of AD on CIZ1-Xi assemblies

The multivalent nature of CIZ1’s interaction with RNA, and the requirement for both domains for the assembly of CIZ1 into SMACs at the Xi^10^, lead us to hypothesize that fragments of CIZ1 encoding only one of its RNA interaction interfaces might have a destabilizing effect. Moreover, based on what we know of CIZ1 genetic deletion and the co-dependency of CIZ1 and *Xist* ^6–8, 24^, we hypothesized that interference with CIZ1 assemblies at Xi would affect Xi chromatin. We modelled this in short-term (one cell cycle) transfection experiments after ectopic expression of GFP-tagged C-terminal protein fragments (Fig. 3A) in murine cells.

**Figure 3.**
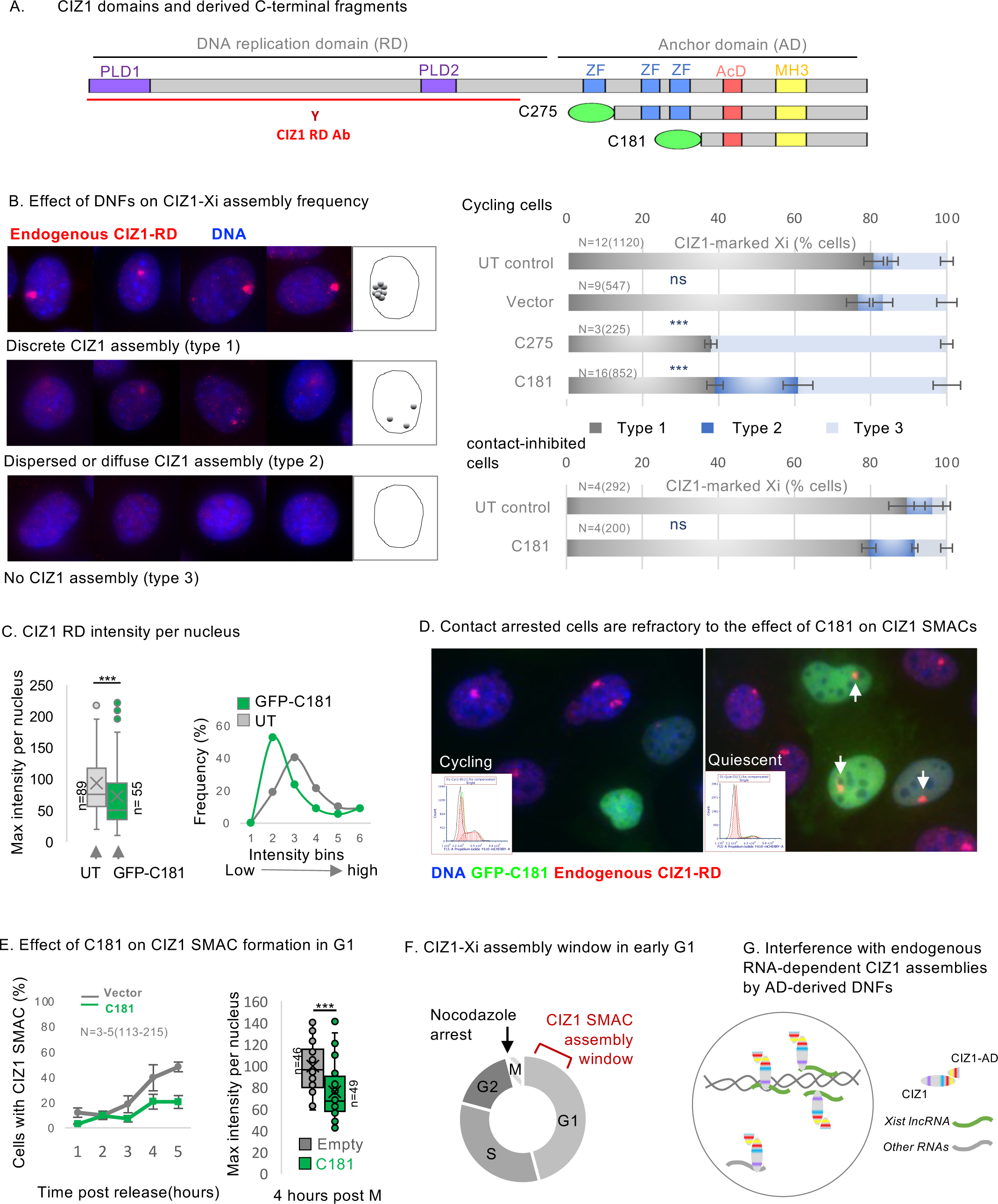
Dispersal of endogenous CIZ1 Xi SMACs by ectopic CIZ1 anchor domain. A) Diagram of murine CIZ1 full-length protein showing prion like domains (PLD)^10^, zinc fingers (ZF), acidic domain (AcD), and Matrin 3 homology domain (MH3, see also SFig.1A). Below, C-terminal protein fragments C275 and C181 correspond to the terminal 275 and 181 amino-acids respectively, both bearing an N terminal GFP tag. CIZ1 RD antibody (red) was raised against a fragment of CIZ1 outside of C275 ^19^, and does not detect the transgene. B) Left, example images of CIZ1-Xi assemblies in D3T3 cells, showing three categories of nuclei with either large discrete CIZ1 SMACs (type 1, upper), no detectable CIZ1 SMAC (type 3, lower), or intermediate assemblies (type 2, middle) which include those that are either dispersed into multiple smaller foci, or diminished in overall size or intensity. CIZ1 is red, DNA is blue. Right, frequency of cells with type 1, 2 or 3 CIZ1 Xi assemblies in untransfected (UT) populations, compared to those expressing empty GFP vector, C275 or C181. N is replicate analyses, with total nuclei inspected shown in parentheses. Error bars show SEM. For cycling cells (upper), no difference in frequencies were observed between untransfected (UT) and empty vector cells (type 1 p=0.72, type 2 p=0.19, type 3 p=0.75), but dispersal was observed in C181 expressing cells compared to UT (type 1 p=1.4×10^-5^, type 2 p=0.0036, type 3 p=0.00032). For contacted cells (lower), none or limited differences in frequencies were observed between UT and C181 expressing cells (type 1 p=0.098, type 2 p=0.34, type 3 p=0.043). All comparisons of replicate analyses are by unpaired t-test. C) CIZ1-RD fluorescence intensity per nucleus in WT female PEFs. Left, box and whisker plot showing mean, median and interquartile range in untransfected (UT) and C181 transfected cells, where n is nuclei measured. Mann-Whitney U test p=0.00087. Right, frequency distribution of intensity values showing a fall in endogenous CIZ1 intensity in cells expressing C181 (green). D) Field images showing untransfected and transfected cycling and contact inhibited D3T3 cells, illustrating effect on endogenous CIZ1 status at Xi (red). Arrows point to resistant CIZ1 Xi assemblies in contact inhibited cells. Inset, flow cytometry profiles of populations stained with propidium iodide, illustrating G1/G0 enrichment in the contacted cell population. E) Impaired reformation of CIZ1 SMACs after release from arrest in M phase in D3T3 cells transduced with C181 compared to vector control. Left, CIZ1-Xi assembly frequency 1-5 hours post release. Right, box and whisker plot showing normalised fluorescence intensity per nucleus at 4 hours, where n indicates the number of nuclei measured in each group. Mann-Whitney U test, p=6.1×10^-6^. F) Illustration showing time window of CIZ1 Xi SMAC assembly early in G1 phase and the point of cell cycle arrest after exposure to nocodazole. G) Illustration showing truncated CIZ1-RNA networks in the presence of C-terminal dominant negative CIZ1 fragments.

Endogenous CIZ1-Xi assemblies were categorised into three phenotypes; cells with a discrete normal assembly, cells with no assembly, or cells with intermediate, dispersed or diminished assemblies (Fig.3B). The C-terminal 275 amino-acids of murine CIZ1 (from exon 10 onwards), here referred to as C275, caused loss or reduction in normal (type 1) assemblies, but did not itself accumulate at Xi. In untransfected cells in the same populations or parallel populations expressing empty GFP vector, CIZ1-Xi assemblies were unaffected, all evidenced via detection of CIZ1 RD epitope (not present in C275). Deletion of two zinc fingers, to produce the smaller C181 fragment, did not abolish the disruptive effect on assembly frequency (Fig.3B) or fluorescence intensity (Fig.3C). Thus, C-terminal fragments of CIZ1 do have the capacity to interfere with endogenous CIZ1-Xi assemblies, and are referred to here as CIZ1 DNFs (dominant-negative fragments).

### CIZ1 assembly dispersal is cell cycle dependent

Not all cells expressing CIZ1 DNFs are depleted of endogenous CIZ1-Xi assemblies. At 24 hours, typically 30-40% remain refractory (Fig.3B), and in those that respond the extent of dispersal is variable. We tested whether cell cycle stage contributes to the heterogenous response, initially by testing contact-inhibited (arrested) cells (Fig.3D). Under these conditions CIZ1-Xi assemblies were refractory to the dominant negative effects of C181 (Fig.3B), while DNFs themselves began to accumulate at sites of endogenous CIZ1 assemblies (SFig.3A), possibly reflecting an extended period of stability. These observations suggest that passage through the cell cycle is required to expose assemblies to a window in which DNFs can exert their effect.

Normally around 80% of female cells (cycling, mouse or human, primary or established non-cancer lines) contain a discrete compact CIZ1-Xi assembly. Since we know that, like *Xist* ^25^, CIZ1-Xi assemblies are lost in mitosis ^7^ we postulated that those cells in which they are not evident have yet to rebuild them and are in early G1 phase. We confirmed this in cells synchronised in mitosis using nocodazole, and found that maximal CIZ1-Xi assembly frequency was reached by four hours after mitotic exit (SFig.3B). Expression of C181 significantly delayed SMAC re-formation during this window, and those that did form had reduced CIZ1 fluorescence intensity (Fig.3E). Thus, the dispersive effect of CIZ1 DNFs is potent during the SMAC assembly window in early G1 phase (Fig.3F,G).

### Role of the MH3 homology domain

To refine the sequence requirements for SMAC dispersal by DNF’s we evaluated a set of six deletion constructs, based on C181 (SFig.3C). All fragments were incorporated into detergent-resistant nuclear structures (SFig.3D), and retained similar capability to interfere with endogenous CIZ1-Xi SMACs, with the exception of one. C181 lacking the Matrin 3 homology domain (ΔMH3) had a small but consistent reduction in potency based on SMAC frequency (SFig.3E), confirmed by measuring fluorescence intensity (SFig.3F). This implicates the MH3 CIZ1:CIZ1 dimerisation interface ^26^ in integrity of endogenous CIZ1 SMACs.

### Consequences of dispersal of CIZ1-Xi assemblies on Xi chromatin

We postulated that dispersal of CIZ1-Xi assemblies by DNFs might mimic the effect on Xi chromatin seen in genetically CIZ1 null primary embryonic fibroblasts (PEFs). In these cells H3K27me3 and H2AK119ub1 are both depleted, and control over PRC target genes, both X-linked genes and elsewhere in the genome, is relaxed ^9^. In single cell cycle experiments, in two cell types, C181 caused a marked reduction in H2AK119ub1-enriched Xi’s but did not affect H3K27me3 (SFig.4A,B.C), while in longer term experiments using lentiviral transduction of C181 (Fig. 4A) both H3K27me3 and H2AK119ub1 were depleted, whether quantified by enriched Xi frequency, or by fluorescence intensity (Fig.4B). Survival of H3K27me3 under conditions where H2AK119ub1 is depleted is consistent with replication-linked dilution of H3K27me3 ^27, 28,9^. Together these data show that DNFs impact histone PTMs.

**Figure 4.**
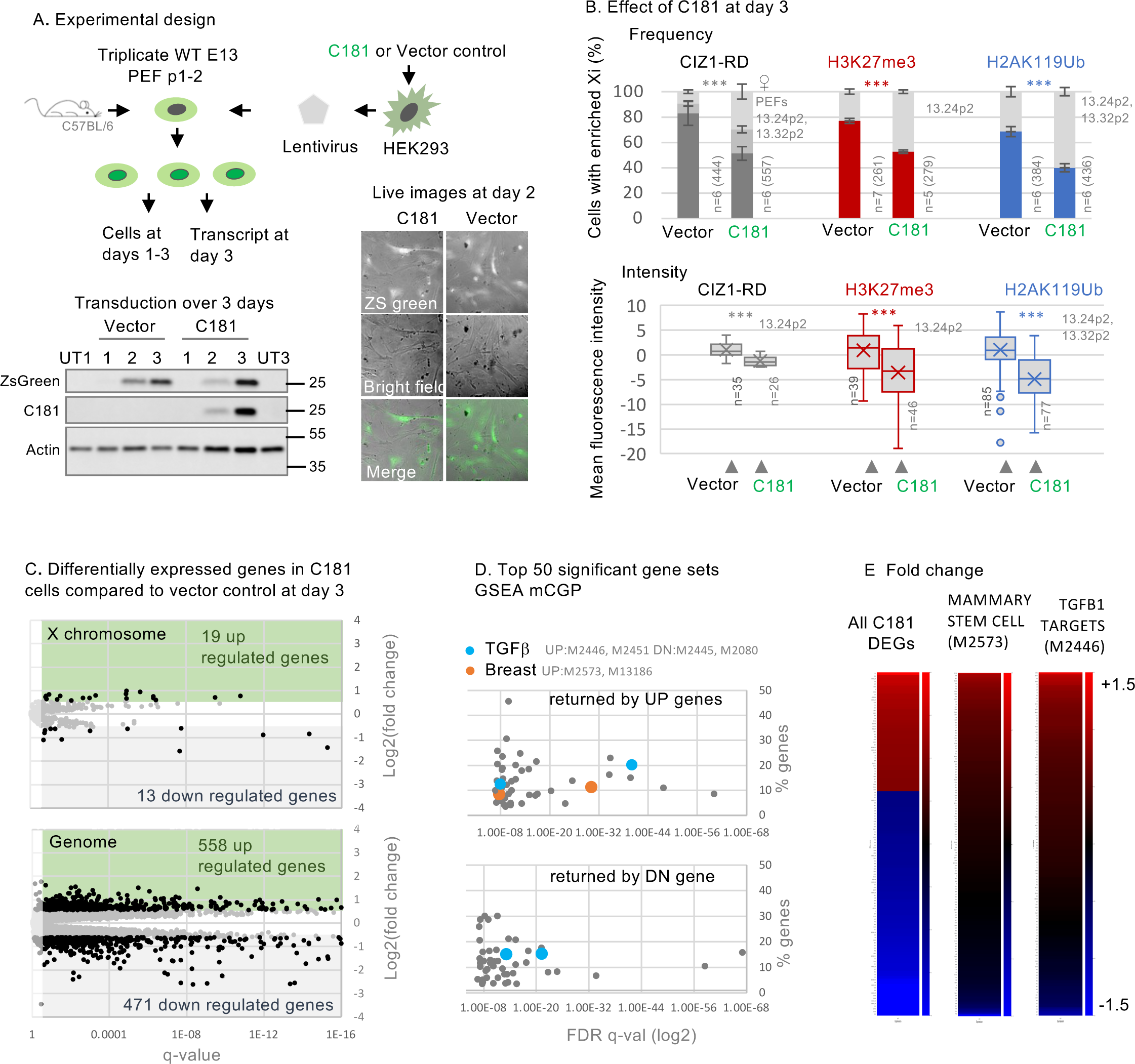
Effect of CIZ1 anchor domain on histones and gene expression. A) Lentivirus encoding C181 and/or ZSGreen was used to infect three independent populations of WT murine primary embryonic fibroblasts (PEFs) at passage 1-2. Below, expression was verified by western blot of ectopic CIZ1 (exon 17) and beta-actin in whole cell lysates over three days, compared to untreated control populations (UT) at days 1 and 3. Below right, live cell images of ZsGreen overlaid on brightfield images of PEFs at day 2 post transduction. B) Comparison of vector only populations to those transduced with C181 showing frequency of cells with CIZ1-Xi assemblies (grey), H3K27me3 (red) or H2AK119ub1 (blue). n denotes replicate analyses with total nuclei inspected in parentheses. PEF cell populations are in grey. Comparisons are by unpaired t-test, where p <0.001 in all cases. Error bars show SEM. Below, box and whisker plots showing mean nuclear intensity measures for cells transduced with C181 or vector control, normalised to the mean of vector only control cells. C) Differentially expressed genes in C181 expressing PEFs, compared to vector control, showing log2 fold change in FPKM against false-detection rate (FDR) corrected q value, and inclusion threshold of q <0.05 and absolute log2FC>1. D) Gene set enrichment analysis of all gene sets derived from chemical or genetic perturbation of murine cells (mCGP). Significance indicator is plotted against % genes in overlap for the top 50 sets returned by C181-induced UP genes, and C181-induced DN genes. Those linked with TGFβ or breast cells are highlighted in blue and red respectively, and set identifiers are given in grey. Source data is given in Supplemental data set 3. E) Heat maps showing C181 DEGs (left, q <0.05 log2FC 1), all genes in mammary stem cell set M2573 ^31^ (middle) and all genes in TGFβ target set M2446 ^59^ (right), where C181-induced fold change of +1.5 or over is maximally red, and less than −1.5 is maximally blue.

The presence of H3K27me3 and H2AK119ub1 is sometimes taken as evidence that PRCs are recruited by lncRNAs causing their enrichment at specific locations, when in fact their residency is often only implied. In some contexts, the balance between addition and removal of marks could be affected in other ways. In our experiments disruption of CIZ1-Xi assemblies by DNFs could deplete H2AK119ub1 in Xi chromatin by reducing recruitment of PRC1, or conversely by deprotecting chromatin and allowing access to de-ubiquitinating enzymes. BAP1 is the catalytic subunit of the deubiquitinating enzyme (DUB) that removes H2AK119ub1, acting to restrict its deposition to specific sites^29^. To begin to distinguish between recruitment and protective functions (SFig.4D) we used PR-619, a broad-spectrum reversible inhibitor of DUBs, including the PR-DUB BAP1^30^. In one cell cycle experiments the immediate (within 24 hours) loss of H2AK119ub1 was significantly blocked by PR-619 (SFig.4B), and in longer (3 day) transduction experiments the same trend was observed (SFig.4E). Moreover, even in genetically CIZ1 null primary cells, in which H2AK119ub1 is absent from Xi chromatin ^9^, its enrichment (but not that of H3K27me3) was restored within 24 hours of exposure to PR-619 (SFig.4F). Thus, loss of CIZ1-Xi assemblies, whether by genetic deletion or dispersal by DNFs, supresses accumulation of H2AK119ub1 in Xi chromatin in a manner dependent on DUB activity. We suggest therefore that CIZ1 assemblies perform a shield function that can protect chromatin from enzymatic attack.

### Effect on gene expression

To confirm that CIZ1 DNFs have the potential to affect gene expression, we analysed transcriptomes of three independent populations of PEFs, transduced with C181 or empty lentiviral vector for 3 days (Fig.4A, SFig.5A). To de-focus analysis from Xi we used primary cells isolated from two female and one male murine embryo. This returned expression changes across all chromosomes (Fig.4C, SFig.5B), including 471 down-regulated genes (DN) and 558 up-regulated genes (UP, FDR q<0.05, log2FC>1, Supplemental Data set 3). Gene set enrichment analysis with those that are named coding genes returned highly significant molecular signatures derived by chemical or genetic perturbation in murine cells (GSEA MSig. mCGP), including sets linked with the developmental regulator TGFβ and mammary stem cell phenotype (SFig.5C). Looking separately at UP and DN genes, sets related to developing breast tissue and mammary stem cell phenotype are returned primarily by UP genes (Fig.4D). Focusing on mammary stem cell phenotype set M2573 ^31^ 25% of all genes in the set are significantly changed by expression of C181 (FDR q<0.05, Fig.4E), and 75% of those are UP. This shows that, similar to germ-line deletion of CIZ1^7^, interference with CIZ1 assemblies in an acute setting can significantly alter gene expression across the genome (SFig.5D), including genes linked with cellular plasticity and cancer. Moreover, as in our previous experiments where CIZ1 is reintroduced against a CIZ1 null background^7^, the effect is rapid (within days) and coincident with changes to epigenetic landscape. Together the data argue for a potent and rapid effect of CIZ1 DNFs, that can change established patterns of gene expression.

**Fig. 5.**
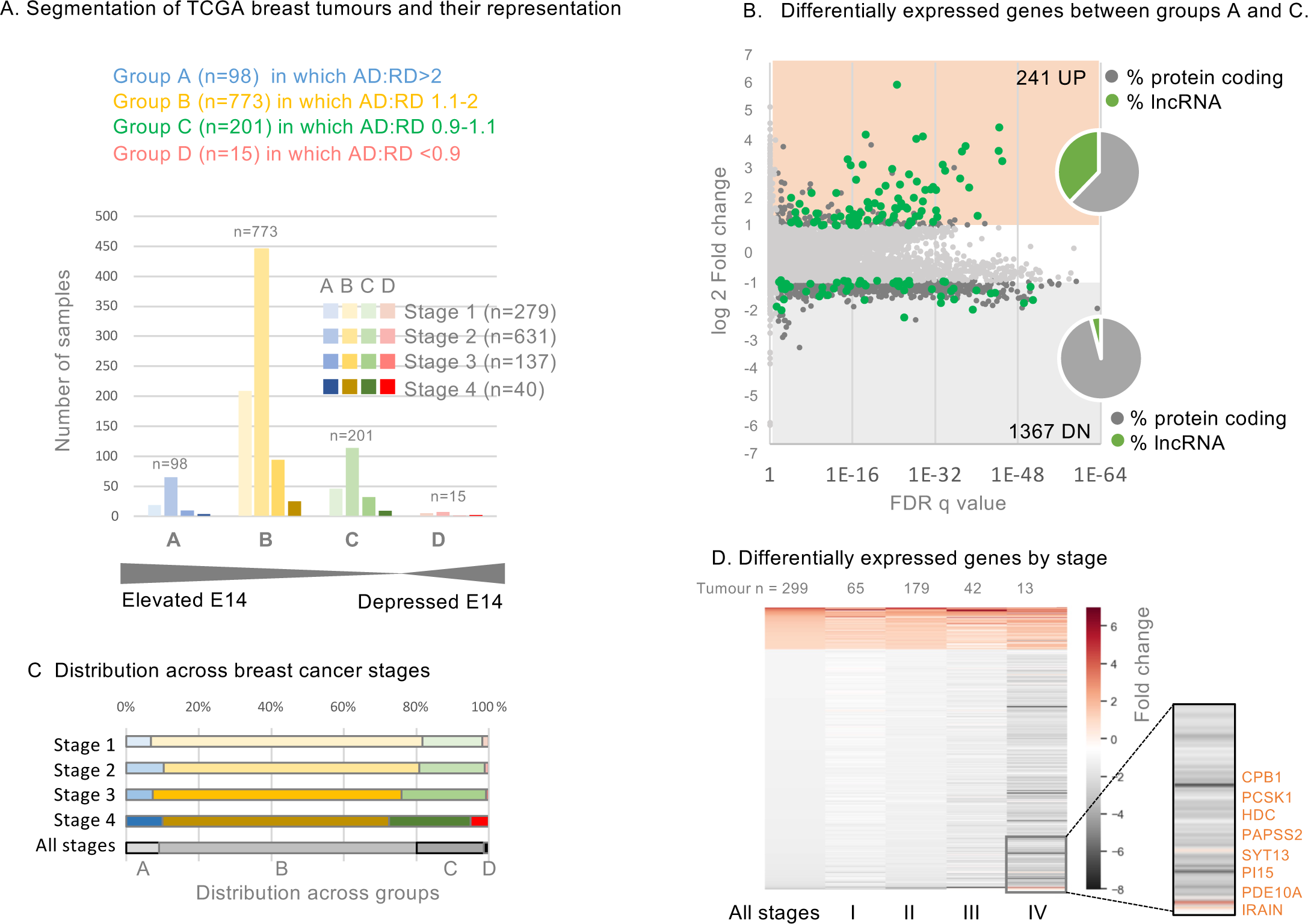
Gene expression in TCGA breast cancers with elevated CIZ1 anchor domain. A) Classification of TCGA breast tumour transcriptomes based on the ratio of CIZ1 AD (exon14) to RD (exon 5), to create a DNF index comprising groups A-D, below shown after segregation by tumours stage. See also Supplemental Dataset 4. B) Differentially expressed genes between groups A (>two-fold elevation of AD) and C (equal +/-10%), showing 241 UP (log2FC≥1) and 1367 DN (log2FC≤1), where q<0.05. Inset, pie charts show the proportion that are lncRNAs (green) or protein coding genes (grey). C) TCGA breast cancers subdivided by stage, showing representation across the DNF index as % (see also SFig.6). D) Heat map showing all differentially expressed genes derived from comparison of groups A and D, and their representation across stages I-IV. Inset, highlight of a small subset of mostly enzyme encoding genes whose expression is suppressed in early stages but which switch to UP genes in stage IV disease.

### Gene expression in human breast cancers

To test whether the disruption of gene expression observed in DNF modelling experiments might be at play in primary human breast cancers, we segmented TCGA breast cancer transcriptomes into four groups A-D (Fig.5A, Supplemental Data set 4), based on the extent of elevation of AD over RD. Gene expression in group A tumours in which exon 14:5 ratio is greater than 2, compared to control group C (where RD and AD are within 10% of even), revealed a massive difference in their transcriptomes (1126 differentially expressed genes FDR q<0.05, log2FC>1, Fig.5B, Supplemental Data set 5).

No significant differences in the proportion of tumours in groups A-D were evident across breast cancer subtypes, or ER/PR/HER2 receptor status subsets^32^ (SFig.6A-C). Similarly, across tumour stages I, II, and III (all subtypes) group A-D profile is close to the cohort profile (Fig.5C, SFig.6D), but shifts at stage IV where a greater proportion are group D (AD:RD ratio favours RD). This mirrors a trend observed in stage IV lung, thyroid and kidney tumours by PCR (SFig.2B) in which RD is more likely to exceed AD. In both contexts however, sample size is too low to draw strong conclusions.

**Fig. 6.**
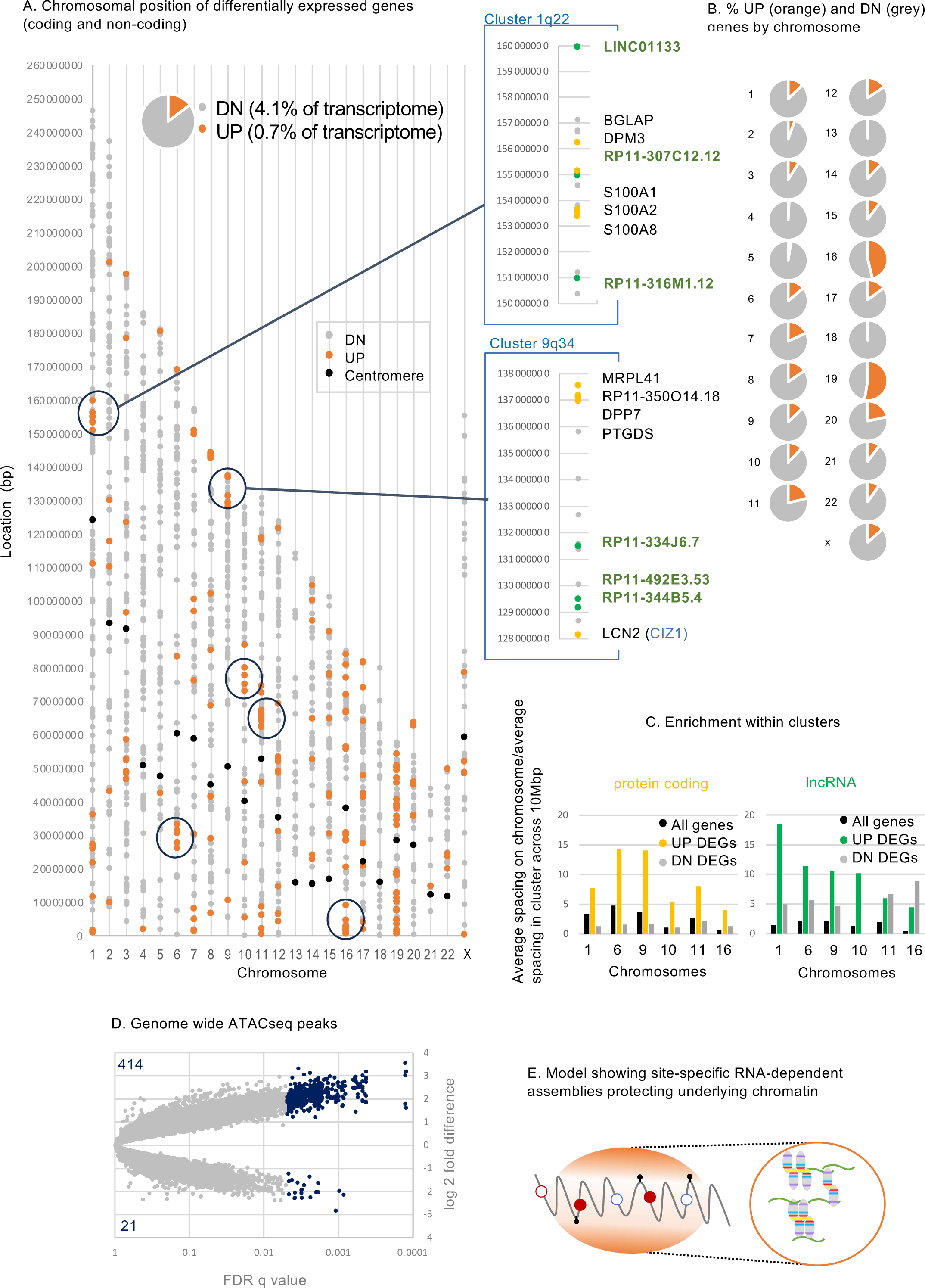
Affected gene clusters, lncRNA enrichment and chromatin accessibility. A) Chromosomal locations of differentially expressed genes derived from comparison of TGGA CIZ1 groups A and D (UP orange, DN grey, q<0.05). Centromere positions in black. Circled clusters are also shown in SFig.7. Right, two example cluster regions on chromosomes 1 (q22), and 19 (q13.2). LncRNAs are green, and protein coding genes yellow. The CIZ1 locus itself is within the circled UP cluster at 9q34. B) Pie charts show the proportion of UP and DN genes by chromosome, highlighting complete absence of UP genes on 18 and high representation on 19. C) Fold gene enrichment in the indicated 10Mb clusters compared to chromosomal average for genes (black), and those that are UP (yellow/green) or DN (grey). Protein coding (left) and lncRNAs (right) are shown separately. D) Genome-wide ATACseq differences between group A and group C TCGA breast tumours, showing all intervals in grey and differently accessible intervals in blue (absolute log2 FC≥1, q<0.05). Over 400 sites are significantly more exposed, compared to 21 that are less exposed. E) Illustration showing localized CIZ1-RNA assemblies surrounding and modulating access to, underlying chromatin.

The differentially expressed genes between groups A and C behave remarkably similarly across stages I, II, and III (Fig.5D), but at stage IV a minority (predominantly enzymes) switch from DN to UP (Fig.5D, segment). Overall, the main conclusion to be drawn from this analysis relates to early-stage disease. Not only is C-terminal elevation evident very early in the course of disease (Fig.2), its effects are also felt early (stage 1), and those effects persist through to later stages.

### Affected chromosomal domains

Of the 1126 genes that are differentially expressed when CIZ1 AD is overrepresented, 15% are UP and 85% are DN. When analysed by location the DN genes are distributed more uniformly than the UP genes, which are clustered (Fig.6A), entirely absent from chromosome 18 and over-represented on gene-dense chromosome 19^33^ (Fig.6B). For six gene clusters of 10Mbp in length (Fig.6C, and circled in 6A, SFig.7A-F) UP regulated protein coding genes are 4-14x denser than the chromosomal average but also enriched 2-6x greater than expected for local gene density. In contrast, the frequency of DN regulated genes reflects local gene density (SFig.7G). This spatially-concentrated UP-regulation is consistent with a CIZ1-related mechanism that normally represses gene expression across large chromosomal domains.

For the six UP gene clusters we asked whether syntenic regions were similarly affected in our mouse model. In fact, all were among those regions encoding UP genes in mouse cultured fibroblasts expressing ectopic AD (SFig.5D). Thus, despite differences in species and cell type, and duration and quantity of AD expression, similarities are observed, arguing for a degree of mechanistic conservation.

Notably, among UP genes 38% encode lncRNAs, compared to only 4% of DN genes (Fig.5B). These are concentrated within clusters of UP-regulated protein coding genes at a density greatly in excess of expected (Fig. 6C, SFig.7G), pointing to a relationship between CIZ1 and lncRNA expression.

Interestingly, differentially expressed lncRNAs that are concentrated in cluster regions are both UP and DN regulated, suggesting functional specialisation. By analogy with the CIZ1-*Xist* complexes that form at Xi, we suggest that CIZ1 normally sequesters lncRNA molecules into RNA-protein assemblies (protecting some), which then modulate access to the locus as a whole. Excess CIZ1 AD expression would be expected to dissolve the assembly and so release the locus.

Exposure of underlying chromatin by DNF-mediated assembly dissolution alters access by deubiquitylases, and might therefore be expected to increase susceptibility to transposases. ATACseq has been performed for a subset of TCGA tumours in group A (n=8) and group C (n=15) to reveal chromatin accessibility across the genome. Using stringent criteria (log2FC>1or <-1, FDR q<0.05) over 400 sites are significantly more exposed in group A than group C (Fig.6D), showing that elevated AD is associated with chromatin accessibility. Exposed sites are located within cluster regions but are also evident in locations that do not host differentially expressed genes (SFig.7A-F, Supplemental Data set 6).

### Affected genes

The many genes (4% of the transcriptome) whose expression is *reduced* in tumours with elevated AD are distributed across all chromosomes, are not enriched in lncRNAs, and the mean fold change is overall less than UP genes (−1.19 compared to +1.58). Together this suggests a different mechanism is at play to that affecting UP genes, and at present it is difficult to form a strong hypothesis about the process. Alone, they return highly significant enrichment scores for gene sets linked with the cellular response to DNA damaging agents (SFig.6E), and when combined with the UP gene set their over 5-fold higher abundance dominates the results.

In contrast, the UP set of 240 spatially regulated genes were highly enriched in breast cancer-related curated gene sets (six of the top 20 significant overlaps FDR q<0.05, SFig.6F), despite cell type signatures identifying primarily lung tissue of fetal origin (SFig.6G). The UP genes also returned five sets associated with other types of cancer, one describing genes under the regulation of EZH2 (catalytic subunit of PRC2 responsible for H3K27me3), and one describing mammary stem cells. Together these studies support the conclusion that excess expression of CIZ1 AD promotes the expression of genes linked with breast cancer.

Notably expression of the *CIZ1* gene itself is not returned as UP or DN regulated, despite the very different domain expression on which groups C and A were defined. This highlights an important deficiency in the way gene expression analysis is typically carried out, with amalgamation of all transcripts for a given gene into one indicator. For CIZ1, the common alteration observed here in breast cancers is not evident from overall expression level data, so it has not yet been captured by large-scale transcriptome studies. Furthermore, there are no recurrent polymorphisms in CIZ1 in adult cancers across 46014 unique samples in COSMIC^34^, so despite apparently profound effects on breast cancer gene expression, CIZ1 is not yet recognised as a ‘cancer’ gene.

## Discussion

The purpose of heterochromatin formation during development is to protect and reinforce cell fate decisions by restricting access to genes. Thus, potential stabilizers of chromatin state, whose mis-expression may lead to heterochromatin instability, are important to understand in relation to degeneration of cellular identity, human disease and aging. Our data suggest that RNA-dependent CIZ1 assemblies, exemplified by the *Xist*- and PLD-dependent CIZ1 SMACs that surround the inactive X chromosome in differentiated cells, normally act as a molecular shield that helps protect Xi heterochromatin from the action of PR-DUBs, and possibly other enzymatic modifiers. ‘Molecular shield’ is one of eight functional classes proposed for phase separating proteins outlined by PhaSePro ^35^. Defined as membrane-less organelles that inactivate reactions by sequestering some of the required components while keeping others outside, a CIZ1 shield would sequester chromatin while excluding PR-DUBs. Questions remain about the structure and influence of such a shield, and whether some molecules penetrate more freely than others.

### Shield loss

We have exploited the easily visualised Xi-associated CIZ1 assemblies as an indicator of dysfunction in breast cancer cells, and as a read out on the solubilizing action of CIZ1 DNFs in a murine model system. Experimentally, exclusion of the N-terminal RD domain which encodes the two PLDs that confer ability to coalesce inside the nucleus ^10^, converts CIZ1 from a SMAC participant into a molecule with the ability to disperse SMACs – a SMAC buster. A shift towards SMAC buster expression is suggested to interfere with normal CIZ1 function in heterochromatin protection, and so contribute to epigenetic deprogramming. Importantly, both SMAC buster sequence elevation in breast cancer cells and experimental DNF transgenes alter the transcriptome and, like deletion of CIZ1^7^, effects are felt across the nucleus, with X-linked genes and other chromosomes similarly affected. Thus, while Xi-associated CIZ1 SMACs offer an important model for visual studies, smaller assemblies associated with other chromosomes are likely also disrupted.

The lack of difference in *Xist* expression between breast cancers with and without elevated AD, and lack of enrichment of differentially expressed genes on the X chromosome has a number of possible explanations: i) lack of homogeneity in response between active and inactive X chromosomes leading to failure to meet the significance thresholds, ii) cancer associated changes that are independent of CIZ1 expression, or iii) lack of Xi sensitivity to loss of CIZ1 assemblies possibly buffered by other repressive mechanisms.

### Susceptible loci

Discrete chromosomal domains are susceptible to SMAC busters at the transcript level, suggesting that the protective effect of CIZ1 assemblies is spatially restricted but broad, extending over domains in excess of 10Mb. Within affected domains, protein coding genes are over-represented but also disproportionately UP regulated, implying both domain-wide de-repression and concentration of genes within CIZ1-protected clusters.

The behaviour of lncRNAs within the same domains is not consistent. While lncRNAs genes are also enriched, and also more likely to be affected than the chromosomal average, this can be UP or DN. Their heterogenous relationship with AD elevation could reflect more than one mechanism. While UP genes may be subject to the same locus de-repression as protein coding genes, the role of lncRNAs in the formation of spatial compartments in the nucleus ^36^ suggest that others might experience transcript preservation upon incorporation into stable locus-specific RNA-protein SMACs.

Taken together these data argue that the chromatin deprotection observed in DNF modelling experiments is at play in breast cancers, and influencing gene expression within specific chromosomal domains, possibly by locally altering the balance between ubiquitination of H2AK119 by PRC1 and its removal by PR-DUBs. Crucially, domain deprotection is evident in early-stage cancers but also persists in later stages, raising the possibility that it is a predisposing influence involved in cancer aetiology. At present the question of what drives DNF expression in early-stage breast cancers is unanswered. Lack of mutations in CIZ1 raises the possibility that DNF expression is itself controlled primarily epigenetically, and that a normal biological context is yet to be found. If DNFs normally confer fluidity on SMACs, for example as cells pass through natural transition states, delays imposed by extrinsic conditions might prolong residency and exposure to the destabilizing effect of DNFs.

### Epigenetic origins of cancer

There remain fundamental questions about the relationship between genetic and epigenetic models of cancer and the question of which comes first is likely to have a range of context specific answers. Mutations in chromatin proteins and their modifiers occur in approximately half of all tumours ^37^ implying that epigenetic instability is a consequence of mutation, yet for some types of tumour no genetic driver mutations are detected ^38^. In fact, it has been shown convincingly that transient depletion of polycomb proteins during Drosophila larval development is sufficient to initiate cancer phenotypes without genetic change ^3^. Our proposal is that expression of CIZ1 DNFs drives disruption of chromatin state in the early stages of tumour development, possibly before acquisition of driver mutations, and certainly before widespread genetic instability. While the TCGA breast cancer analysis suggests this, direct modelling of the impact of DNFs by introduction to normal cells shows unequivocally their ability to drive widespread changes in gene expression.

## Author contributions

GT, EL, ES, JA, AM, DC designed experiments. GT, EL, ES, HC, RG, AA, AM, DC performed experiments. GT, JA and DC wrote the paper.

## Acknowledgments.

We thank Will Brackenbury and Jonathan Godwin for cells, Christian Fermer of FDAB for anti-CIZ1 antibodies, and former colleagues Faisal Abdel Rahman, Jennifer Munkley, Louisa Williamson, Gillian Higgins, Julie Tucker, Karen Clegg and Matt Dowson. We acknowledge the role of the Breast Cancer Now Tissue Bank in collecting and making available the primary normal breast epithelial cells used here, and the patients who donated their tissues. Protein domain analysis was funded by the Georgina Gatenby PhD scholarship to GT, human CIZ1 gene expression array work by Cizzle Biotech, reanalysis of TCGA data by York Against Cancer, and remaining work by MR/V029088 and a Royal Society Leverhulme Trust Fellowship to DC.

## Methods

### Materials availability and contacts

Further information and requests for resources and reagents should be directed to the lead contacts, Gabrielle Turvey and Dawn Coverley.

### Human Primary Cells

Primary human mammary epithelial cells (HMECs) were cultured at 37°C with 5% CO_2_ in MEBM basal medium (Lonza) supplemented with MEGM SingleQuots (Lonza) on culture dishes coated in collagen (Thistle Scientific), and sampled at passages 1-2. HMECs were acquired with informed consent from three donors by Breast Cancer Now Tissue bank under NHS ethical approval, and accessed under local approval from the University of York Department of Biology Research Ethics Committee.

### Human Cell Lines

All cell lines used are of female origin and were authenticated for this study by Eurofins Genomics human cell line authentication service (Eurofins Medigenomix Forensik GmbH), which returned the expected identities with 92-100% confidence in all cases. MCF-10A is a non-tumorigenic epithelial cell line established from human mammary gland with fibrocystic disease. MCF7 is a poorly-aggressive and non-invasive triple receptor positive human breast cancer cell line established from epithelial cells isolated from a metastatic mammary adenocarcinoma. BT-474 is a human breast cancer cell line established from a malignant ductal carcinoma of the breast that overexpresses human epidermal growth factors receptors 2 (HER-2) and oestrogen receptors (ER). SK-BR-3 was established from a malignant adenocarcinoma of the breast that overexpresses HER-2. MDA-MB-231 is a human breast epithelial cancer cell line established from a metastatic poorly differentiated triple-negative mammary adenocarcinoma. Cells were cultured in the following media: MCF-10A, MEGM, 5% horse serum,10 µg/mL hydrocortisone, 20 ng/mL EGF, 500 ng/mL insulin, 100 ng/mL cholera toxin, 1% PSG; MCF7, EMEM, 10% foetal bovine serum (FBS), 1% Penicillin-Streptomycin-Glutamine (PSG) (Gibco); BT-474 and SK-BR-3, DMEM, 10% FBS, 1% PSG; MDA-MB-231, DMEM, 5% FBS, 1% PSG.

### Mouse Primary Cells

All mouse primary embryonic fibroblast (PEF) strains (WT 13.24, 13.31, 13.32, 13.33, 13.27, 45.1fc and CIZ1 null 13.17, 41.2fa) were derived from day 13 embryos from C57BL/6 mice as previously described ^7, 9^. CIZ1 null mice were generated from C57BL/6 ES clone IST13830B6 (TIGM) harbouring a neomycin resistance gene trap inserted downstream of exon 1. The absence of *Ciz1*/CIZ1 in homozygous progeny was confirmed by qPCR, immunofluorescence and immunoblot. Breeding of mice and all work with animal models was carried out under UK Home Office license and with approval of the Animal Welfare and Ethical Review Body at the University of York. PEFs were cultured in 4.5g/L glucose DMEM containing 10% FBS, 1% PSG up to a maximum of passage 3. After passage 4 these cells are referred to as MEFs and were not used here.

### Mouse Cell Line

The female D3T3 cell line was cultured as described ^9^ in DMEM, 10% FBS, 1% PSG (Gibco).

### Site-directed mutagenesis

Mutagenic primers that contain additions, substitutions or deletions of murine CIZ1 by PCR mutagenesis created for this study, are listed in Resource table. All plasmids were sequence verified to confirm mutations (Eurofins TubeSeq Service).

### Transient Transfection

For analysis in cycling cells, cells were seeded on 13mm glass coverslips at approximately 30% confluency one day prior to transfection, to produce populations at ∼60% confluency at time of transfection. Coverslips were transferred to individual wells in 24 well plate in 500μl media prior to transfection. For each coverslip 50 μl Opti-MEM® Medium (Gibco) was mixed with 1.5 μl X2 Transfection Reagent (Mirus) and 200ng plasmid DNA (pEGFP-C2 with or without inserts derived from CIZ1), incubated for 30 min, then applied to cells dropwise. Coverslips were fixed and processed for immunofluorescence typically 24hrs later. For contact inhibited cells, cells were plated across a range of densities by serial dilution two days prior to transfection. Coverslips at greater than 90% confluency were selected for transfection, and processed as above.

### Cell synchrony

D3T3 cells were arrested in mitosis or S phase using 50 ng/ml nocodazole (Sigma) for 16-24 h, or 2.5 mM thymidine (Sigma) for 24 h, respectively. Cells arrested in M phase were isolated by mitotic shake off and re-plated onto glass coverslips for analysis post release. Cells held in S phase grown on glass coverslips were released by washing twice with PBS, then replacing with fresh media. In transduced cell populations, cells were arrested approximately 48 hrs post transduction for 16-24 hr, then released and analysed. To facilitate retention of mitotic cells, coverslips were fixed prior to permeabilization.

### Flow cytometry

Cells were isolated from 9 cm culture plates by trypsinization and resuspended in 100 μl cold PBS to obtain a single cell suspension, then stored at −20°C after addition of 1.5 ml cold 70% ethanol. For analysis cells were pelleted and resuspended in PBS (500,000 cells/mL), and 55 μl 10x FACS mix (1 mg/ml propidium iodide, 4% v/v Triton X-100, 10xPBS) was added per 500 μl of cell suspension. DNA content was measured using CytoFLEX (Beckman Coulter) at excitation 561 nm/emission 610/20 for detection of the DNA binding dye propidium iodide ^39^. A minimum of 5,000 single cells per sample were recorded for analysis using cell cycle algorithm software FCS Express V7 (Dotmatics).

### Inhibitors

To measure the impact of inhibition of PR-DUBs, 5µM PR-619 (Bio-Techne) was applied to PEFs 16 h post transduction for 32 hrs, to collect cells 48 h post transduction. In transient transfection experiments PR-619 was used at 5 µM for 24 h throughout the transfection window.

### Lentivirus transduction

Bicistronic ZsGreen/C181-bearing virus and ZsGreen alone bearing virus were produced in the Lenti-X™ 293T subclone of human embryonic kidney (HEK) cells. 8×10^5^ HEK cells were seeded per well in a 6 well plate prior to transfection with plasmids. For transfection of each well, 1 µg transfer vector, 0.75 µg packaging plasmid, 0.25 µg envelope plasmid diluted in 100 µl optiMEM (Gibco) was mixed with 20 µl of PolyFect transfection reagent (Qiagen) and incubated for 5-10 min at room temperature to allow complex formation. 0.6 ml of cell growth medium was added and gently mixed then transferred to one well. Cells were incubated for 16 h, then media was replaced with fresh growth medium (supplemented with addition of HEPES to a final concentration of 20 mM). At 48 h post-transfection the supernatant containing virus was harvested and filtered through a low-protein binding filter (0.45µm, Sarstedt) to remove HEK debris. Viral supernatant was supplemented with 4 μg/ml polybrene (Sigma), and transferred to recipient PEFs or D3T3 cells. Transduction was monitored by emergence of cytoplasmic ZsGreen, and showed that close to 100% of the cells were transduced after approximately 48h.

### Immunofluorescence

Cells grown on coverslips were washed in cytoskeletal buffer (10 mM PIPES/KOH pH 6.8, 100 mM NaCl, 300 mM sucrose, 1 mM EGTA, 1 mM MgCl_2_) with 0.1% v/v Triton X-100, and fixed in 4% w/v paraformaldehyde (PFA). Where indicated Triton X-100 was left out of CSK (unextracted cells), or additional 400mM NaCl was added (high-salt extraction). After fixation all coverslips were blocked in antibody buffer (AB) (1xPBS, 10 mg/ml BSA, 0.02% w/v SDS, and 0.1% v/v Triton X-100) for 30 min, incubated with primary antibodies for 1 h at 37°C, washed three times with AB, incubated with secondary antibodies for 1 h at 37°C, washed three times with AB, and mounted on glass slides with Vectashield containing DAPI (Vector Labs). Primary antibodies are detailed in Resource table. Anti-human CIZ1 monoclonal antibody 87 was generated by Fujirebio Diagnostic Antibodies (FDAB). Anti-species antibodies (ThermoFisher) labelled with Alexa Fluor 568 (red) or 488 (green) was used for detection in all cases. Fluorescence images were captured using a Zeiss Axiovert 200M fitted with a 63×/1.40 Plan-Apochromat objective and Zeiss filter sets 2, 10, and 15 (G365 FT395 LP420, BP450-490 FT510 BP515-565, and BP546/12 FT580 LP590), using Axiocam 506 mono and Axiovision image acquisition software (SE64 release 4.9.1) through Zeiss Immersol 518F. For each antibody constant image capture parameters were used to generate image sets within an experiment, on which quantitative analysis was performed, in all cases from unmodified raw images.

### Phenotype Scoring in Dispersal Assay

Cells were classified by eye across replicate experiments, and across 2-3 replicate coverslips per condition within an experiment. Avoidance of bias was achieved by verification by independent workers in all cases, and blinded analysis in some cases. In the three-tier scoring system cells were categorised as either having a compact CIZ1 Xi assembly (type 1) or not (type 3), or were assigned to an intermediate category (type 2) in which CIZ1 assemblies were reduced or diffuse, or made up of locally dispersed particles. Examples are shown. Empty vectors (EV) are used as a negative control, and WT-C181 as positive control in experiments to test the effect of mutants. In transient transfection experiments, untransfected (not green) cells within test populations serve as internal controls on each coverslip.

### Image analysis in dispersal assays

For measurement of the effect of CIZ1 fragments on endogenous CIZ1 or histone PTMs, sets of images including test and controls samples, were processed in parallel and imaged with identical parameters in one sitting. All intensity measurements were conducted on unedited, unenhanced raw image sets. FIJI identified regions of interest (ROI) within DAPI stained fields of nuclei using auto-thresholding with Otsu setting to create a binary mask that defined nuclear perimeters. ROI’s were applied to antibody-detected fluorescence image layers, to generate nuclear intensity means, minimum, and maxima per ROI, and area of each nucleus. In female nuclei, to obtain a surrogate estimate of Xi-assembly intensity, overall nuclear maxima were used. Where two or more data sets were combined (for example two C181/vector control pairs) from experiments performed on different days or with different PEF populations, data was amalgamated after normalization of values to the average of the control set in each case. For reproduction, images were digitally enhanced to remove background fluorescence or increase brightness using FIJI. Identical manipulations were applied within an experiment, so that for example, the intensity of staining with and without transfection, or before and after extraction, is accurately represented.

### Quantitative RT-PCR

To quantify relative expression of CIZ1 amplicons in a wide range of primary tumours, TissueScan_TM_ Tumour cDNA arrays from OriGene Technologies, Inc. (Rockville, MD) containing 2-3 ng of cDNA were analysed by qPCR. The RNA was collected under IRB approved protocols, and array details with tumour classification is given in Supplemental Data set 2. Tumour classifications and abstracted pathology reports are given at: http://www.origene.com/qPCR/Tissue-qPCR-Arrays.aspx. cDNA was normalized using β-actin by the supplier, and we used our own amplification of β-actin where indicated. In most cases results for CIZ1 amplicon expression were expressed relative to one another, rather than to another gene, and normalised to control samples so that change in ratio across cancer samples is apparent. Reactions were carried out in 25 µl volumes with 12.5 µl Taqman master mix (Applied Biosystems), 1 µl of each 10 µM primer and 1 µl 10 µM probe. Primers (Sigma Aldrich) and probes (MWG) are specified in Resource table. Primers 9 and 10 were combined with a probe in exon 5 to generate detection tool set DT5, primers 13 and 14 with probe 7 (DT7), primers 1 and 2 with probe 14 (DT14), and primers 6 and 7 with probe 16 (DT16). Primer efficiencies were greater than 90% in all cases, and the relative amplification efficiencies of RD and AD tools were routinely checked using plasmid template with coupled and equal levels of RD and AD. Data was generated using an ABI 7000, SDS v1.2. (Applied Biosystems), using 50°C (2 min), 95°C (10 min), then 50 cycles of 95°C (15 s), 60°C (1 min). Relative expression was calculated using the comparative Ct method using the formula, 2^-ΔΔCt^ ^40^, and results expressed relative to the mean of normal cells or tissue in each array, or to the lowest stage tumour in the array, as indicated.

### Mouse transcriptomics

Primary murine embryonic fibroblasts 13.31, 13.32 (WT female) and 13.33 (WT male) were transduced with virus bearing either the empty pLVX-EF1a-IRES-ZsGreen1 vector (Takara), or the same plasmid expressing the coding sequence of C181 as described in lentivirus transduction. After 72 hrs, RNA was extracted with TRIzol (Ambion 15596-026) following manufacturer’s instructions and RNA pellets were resuspended in nuclease-free water. Isolated RNA was treated with DNase (Roche 04716728001) before quality analysis by agarose gel, NanoDrop spectrophotometer and Agilent 2100 Bioanalyzer. Library preparation and sequencing was undertaken by Biomarker Technologies (BMKGene), using the NEBNext UltraTM RNA Library Prep Kit for Illumina® (NEB, USA), with enrichment for mRNA using oligo(dT)-magnetic beads, followed by random fragmentation of enriched mRNA in fragmentation buffer. cDNA was synthesized using random primers followed by purification with AMPure XP beads, end repair, dA-tailing, adaptor ligation, PCR enrichment and further AMPure XP purification to select fragments within size range of 300-400 bp. Library quality was assessed using the Agilent 2100 Bioanalyzer system and sequenced using an Illumina platform, using paired-end sequencing to generate at least 15.31Gb clean data per sample, with minimum 93.16% of clean data, quality score of Q30. Low quality sequence reads and adaptor sequences were removed and the resulting high-quality reads were aligned to version GRCm38 of the mouse genome using HISAT2 ^41^. Transcriptomes were assembled and gene expression quantified using StringTie ^42^ against the Ensemble release 79 annotation. Differential gene expression analysis was performed by DESeq2 ^43^. Bioinformatic analysis and plots were generated from the average of transcripts per million plus one (TPM+1) values for each treatment condition to exclude very lowly expressed transcripts using Spyder (v.5.3.3) accessed by Anaconda Distribution (v.2.3.2). Scatter plots were generated using the pandas, numpy and matplotlib modules. Volcano plots were generated using the pandas and bioinfokit modules. Principle component analysis (PCA) plots were generated using the pandas, sklearn, seaborn and matplotlib modules. Gene set enrichment analysis was performed using the GSEAPY module, with genes preranked based on generation of a π value ^44^ as calculated by multiplication of the log_2_FC of the average TPM by the -log_10_(q value), and also separately for UP and DN differentially expressed genes.

### Patient and Cell Line Bioinformatics

Aligned RNA sequencing data for 1095 primary breast cancer samples from The Cancer Genome Atlas were accessed under dbGaP project 25297. Secondary metastatic breast samples were excluded from analysis. Data were downloaded using the Genomic Data Commons command line client v1.5.0. FASTQ files were regenerated from sample BAM files using samtools v1.10 ^45^ to exclude secondary and supplementary alignments, and then BEDTools v2.27.1 bamToFastq ^46^. No additional quality control steps were performed on the extracted read files. Reads were aligned to the GRCh38 Gencode primary assembly and to individual CIZ1 transcripts (from Gencode v38 and ^47^) using HISAT2 v2.2.0 ^41^. Reads were also pseudo-aligned to the Gencode v38 full annotation transcriptome file with kallisto v0.46.0 ^48^, quantified and aggregated to gene-level transcripts per million (TPM) expression values using tximport v1.24.0 ^49^. The same expression analysis pipeline was also completed on publicly available RNA sequencing data from four breast cancer cell lines - MCF7 (SRR8615758), BT-474 (SRR8616195), SK-BR-3 (SRR8615677), MDA-MB-231 SRR8615767) and the breast epithelium transformed cell lines MCF-10A (SRR12877369).

### Analysis of domain imbalance

CIZ1 encodes 16 translated exons (2-17), plus at least three alternative untranslated exons 1s which were excluded from the present analysis. CIZ1 exon transcript read coverage in TCGA breast tumours was generated by alignments to both transcript and full genome and inspected manually for novel, well-supported splice junctions in IGV Desktop for Windows v2.8.2 ^49^. Outputs were normalised to the canonical (ENST00000372938.10) exon 7 coverage, and stratified by tumour stage. To develop an index of the degree of CIZ1 AD and RD domain imbalance, the ratio of exon 14 to exon 5 was calculated for individual tumours after mapping to CIZ1 transcript. The 10 stage II patients with most imbalanced AD:RD were used to rule out broader imbalance in 3’ coverage within the libraries. These were TCGA-E9-A54Y, TCGA-LL-A6FR, TCGA-E9-A3X8, TCGA-AQ-A54O, TCGA-AQ-A54N, TCGA-WT-AB41, TCGA-A2-A3XV, TCGA-LL-A5YL, TCGA-GM-A2DB, TCGA-AO-A03N, which were similar across *ESR1* and *TP53* exons. To compare gene expression in TCGA tumours with and without domain imbalance, a sub group A (n=98) in which AD (exon 14):RD (exon5) normalised TPM ratio exceeds 2, and a sub group C (n=201) in which the ratio is within 0.9-1.1 (10% variance) were identified (supplemental dataset 4). Intermediate group B (ratio 1.1-1.99, n=773) and group D (<0.9, n=15) are also shown. Differentially expressed genes (DEGs, absolute log2 fold change >1, FDR q-value 0.05) between groups A and C reveal those whose expression correlates with elevated AD. The representation of groups A-D across breast cancer subtypes was based on histological classifications^32^, and on tumour stage classifications associated with accessed transcriptomes ^32^ (listed in supplemental dataset 4), and compared to the cohort as a whole.

### ATACseq analysis

Aligned ATAC sequencing data for all available sub group A (n=8) and sub group C (n=15) primary breast cancer samples from The Cancer Genome Atlas were accessed under dbGaP project 25297. BAM files were filtered to remove unmapped reads and sorted by read name using SAMtools, before conversion to FASTQ format using BEDTools bamToFastq. FASTQ files were processed and analysed using the nfcore/atacseq v2.1.2 workflow using default parameters. DESeq2 was used to identify differentially accessible regions based on read counts within consensus peaks, stringently filtered for absolute log2 fold change >1, FDR q-value 0.05.

### Positional analysis

Chromosome maps were generated by segregating DEGs by UP or DN then by chromosome, and plotting by the start position of each mapped gene in Excel. Centromere positions were extracted from mouse genome build GRCm39/mm39 and human GRCh38.p14 using USCC genome browser. The plotted murine gene cohort includes 1029 C181 DEGs (FDR q<0.05, log2FC>1 or <-1), and human gene cohort are 1126 DEGs related to 2-fold elevated AD in TCGA patients (FDR q<0.05, log2FC>1 or <-1). Average UP and DN gene density was calculated based on sequenced human chromosome lengths given in NCBI Genes and Disease, and locations enriched for UP genes selected based on increased density across 10Mb domains. For the 6 human clusters analysed, UP DEG enrichment exceeded gene enrichment by 2-12 fold. Syntenic regions in the murine genome for human UP gene clusters were identified using Ensemble Chromosome view synteny tab.

### Quantification and statistical analysis

For analysis of CIZ1 SMAC frequencies, a variable number (>3) of technical replicates and independent counts (N value) were conducted per experiment as indicated, allowing generation of ± SEM. The number of cells scored is stated individually in each experiment (n value). Wherever possible at least two independently isolated PEF lines were used across the experimentation relating to each question. Statistical analysis was carried out using SPSS release 2021 ver.28.0 for Mac (IBM), or Microsoft Excel for Mac ver.16.73.3 Parametric or non-parametric tests were utilised where appropriate; for comparison between two data sets a two-sample unpaired t-test or a Mann-Whitney U test was utilised, for comparison between three or more data sets a one-way ANOVA was followed by an appropriate post-hoc test. Statistical tests used in each analysis are stated in the figure legend, with p value, where asterisks indicate *p< 0.05, **p< 0.01 and ***p<0.001. For qRTPCR data Pearson correlation coefficient was used to compare CIZ1 RD and AD amplicon expression, and linear regression trendlines were applied using Excel. Graphs were generated using Excel.

## Resources

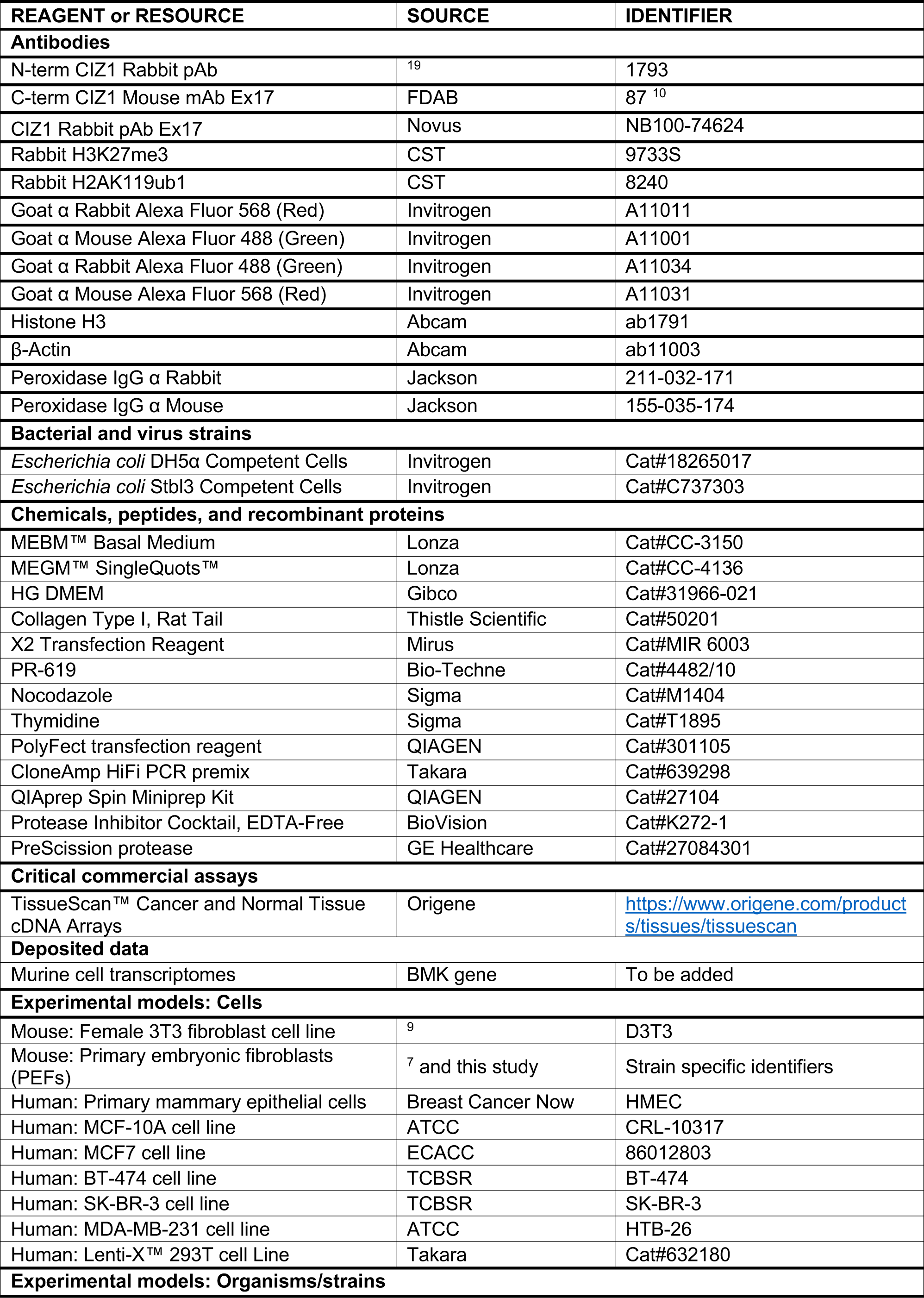
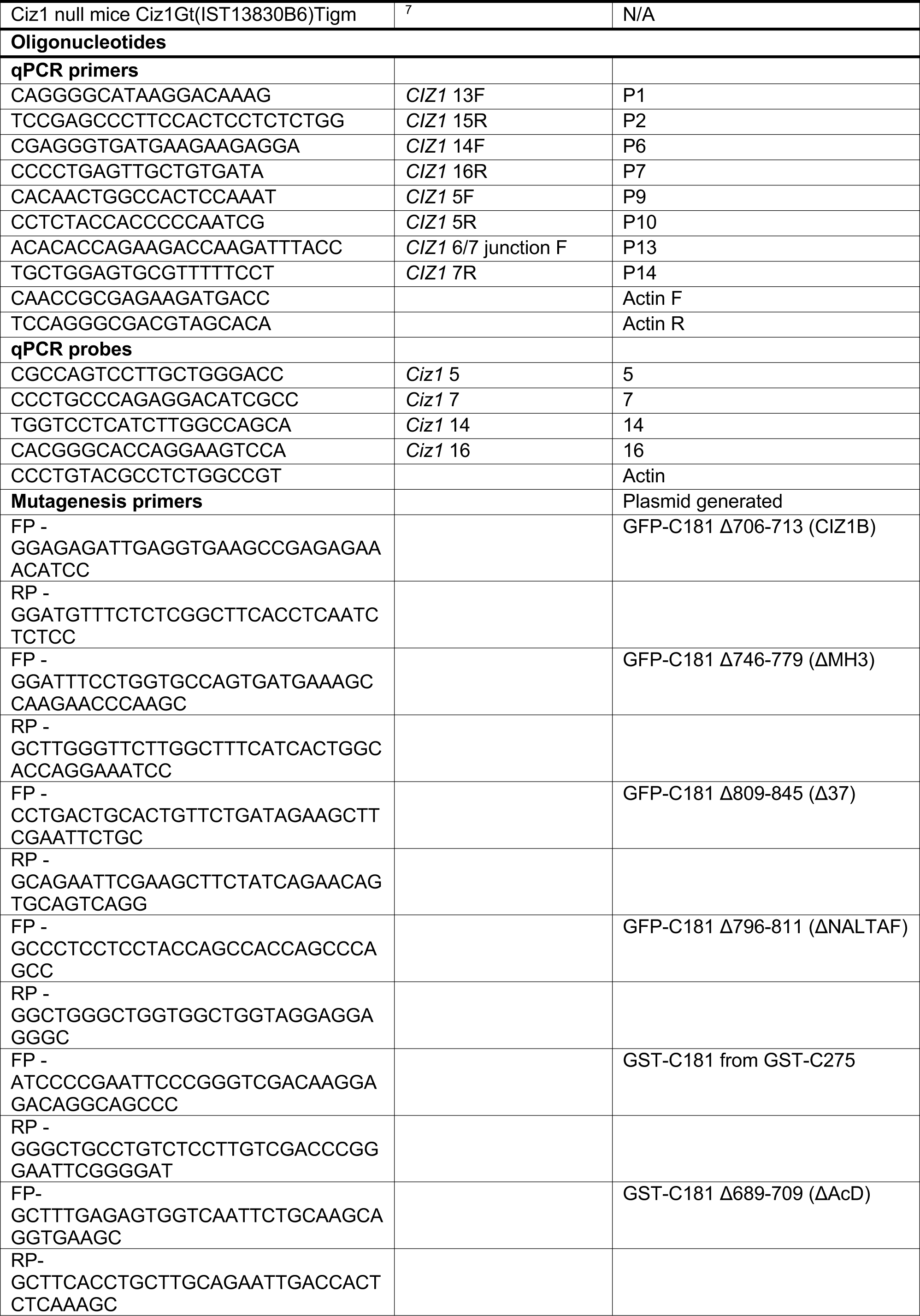
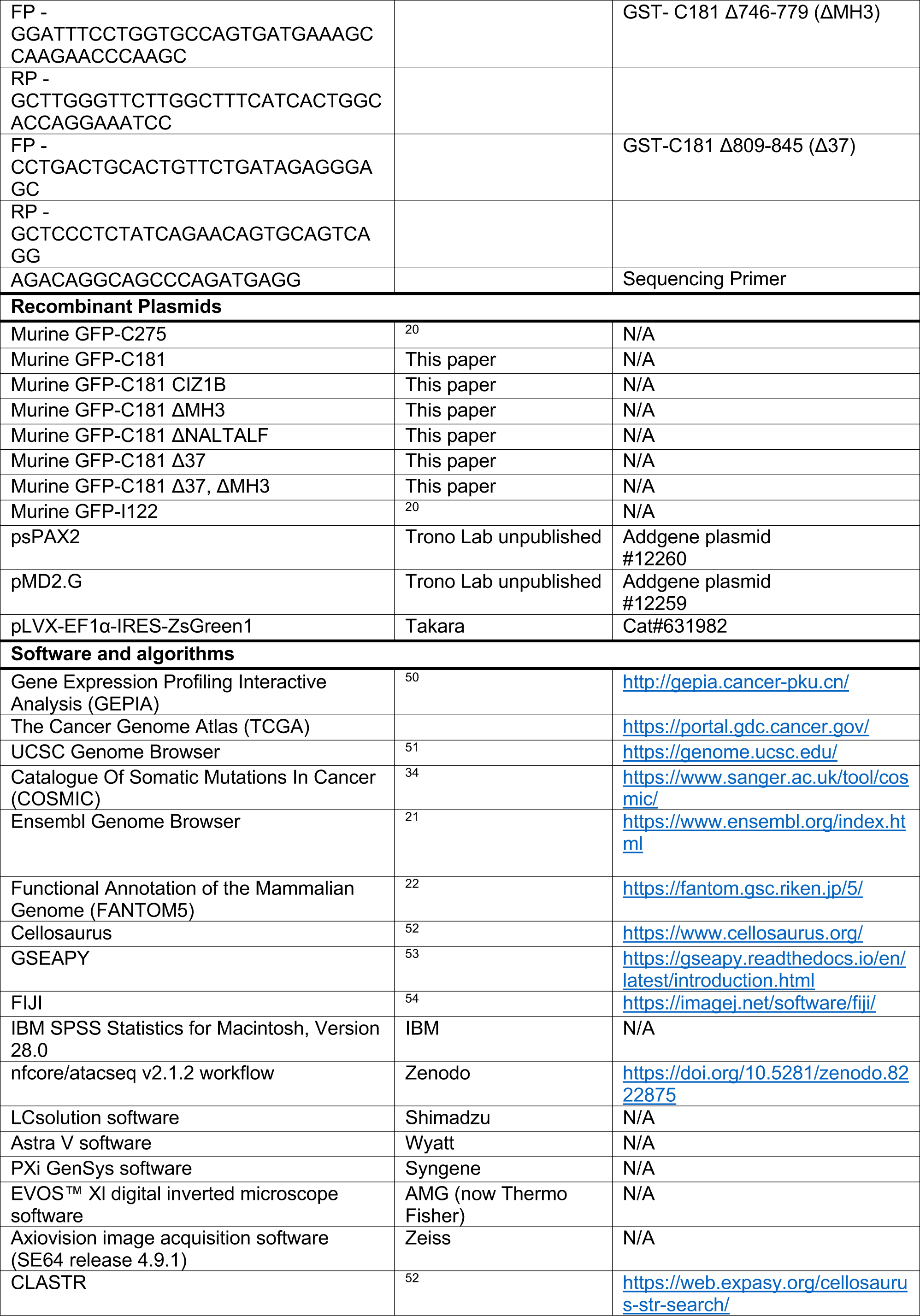
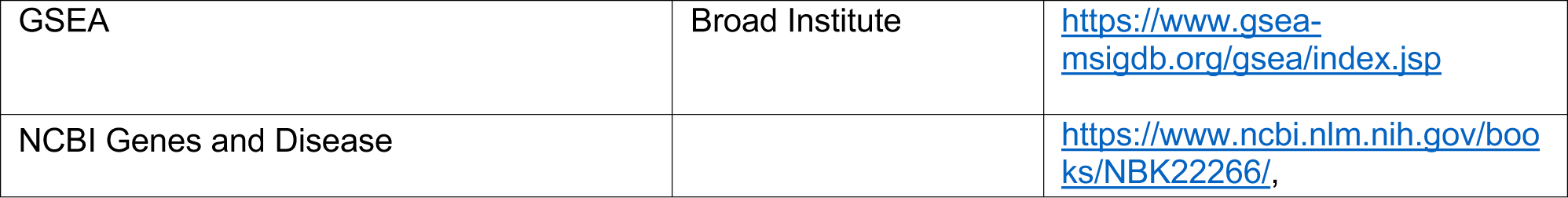

## List of supplemental materials

**Supplemental data set 1** (Xl file)

CIZ1 exon expression (cell lines and tumour summary)

**Supplemental data set 2** (Xl file)

Data and primary tumour designations for QRTPCR array analysis.

**Supplemental data set 3** (Xl file)

DNF effect on murine transcriptomics and GSEA analysis

**Supplemental data set 4** (Xl file)

TCGA BRCA transcriptomics, segmentation and clinical metadata

**Supplemental data set 5** (Xl file)

Gene expression in BRCA group A compared to C

**Supplemental data set 6** (Xl file)

ATACseq in group A compared to C

## References

1. Flavahan, W.A., Gaskell, E. & Bernstein, B.E. Epigenetic plasticity and the hallmarks of cancer. Science 357 (2017).

2. Hanahan, D. Hallmarks of Cancer: New Dimensions. Cancer Discov 12, 31–46 (2022).

3. Parreno, V. et al. Transient loss of Polycomb components induces an epigenetic cancer fate. Nature (2024).

4. Brockdorff, N. et al. The product of the mouse Xist gene is a 15 kb inactive X-specific transcript containing no conserved ORF and located in the nucleus. Cell 71, 515–526 (1992).

5. Brown, C.J. et al. The human XIST gene: analysis of a 17 kb inactive X-specific RNA that contains conserved repeats and is highly localized within the nucleus. Cell 71, 527–542 (1992).

6. Markaki, Y. et al. Xist nucleates local protein gradients to propagate silencing across the X chromosome. Cell 184, 6174–6192 e6132 (2021).

7. Ridings-Figueroa, R. et al. The nuclear matrix protein CIZ1 facilitates localization of Xist RNA to the inactive X-chromosome territory. Genes Dev 31, 876–888 (2017).

8. Sunwoo, H., Colognori, D., Froberg, J.E., Jeon, Y. & Lee, J.T. Repeat E anchors Xist RNA to the inactive X chromosomal compartment through CDKN1A-interacting protein (CIZ1). Proc Natl Acad Sci U S A 114, 10654–10659 (2017).

9. Stewart, E.R. et al. Maintenance of epigenetic landscape requires CIZ1 and is corrupted in differentiated fibroblasts in long-term culture. Nat Commun 10, 460 (2019).

10. Sofi, S. et al. Prion-like domains drive CIZ1 assembly formation at the inactive X chromosome. J Cell Biol 221 (2022).

11. Moore, K.L. & Barr, M.L. The sex chromatin in human malignant tissues. Br J Cancer 11, 384–390 (1957).

12. Chaligne, R. et al. The inactive X chromosome is epigenetically unstable and transcriptionally labile in breast cancer. Genome Res 25, 488–503 (2015).

13. Sirchia, S.M. et al. Misbehaviour of XIST RNA in breast cancer cells. PLoS One 4, e5559 (2009).

14. Batra, R.N. et al. DNA methylation landscapes of 1538 breast cancers reveal a replication-linked clock, epigenomic instability and cis-regulation. Nat Commun 12, 5406 (2021).

15. Locke, W.J. & Clark, S.J. Epigenome remodelling in breast cancer: insights from an early in vitro model of carcinogenesis. Breast Cancer Res 14, 215 (2012).

16. Locke, W.J. et al. Coordinated epigenetic remodelling of transcriptional networks occurs during early breast carcinogenesis. Clin Epigenetics 7, 52 (2015).

17. Dixon-McDougall, T. & Brown, C.J. Multiple distinct domains of human XIST are required to coordinate gene silencing and subsequent heterochromatin formation. Epigenetics Chromatin 15, 6 (2022).

18. Valledor, M. et al. Early chromosome condensation by XIST builds A-repeat RNA density that facilitates gene silencing. Cell Rep 42, 112686 (2023).

19. Coverley, D., Marr, J. & Ainscough, J. Ciz1 promotes mammalian DNA replication. Journal of Cell Science 118, 101–112 (2005).

20. Ainscough, J.F. et al. C-terminal domains deliver the DNA replication factor Ciz1 to the nuclear matrix. Journal of Cell Science 120, 115–124 (2007).

21. Cunningham, F. et al. Ensembl 2022. Nucleic Acids Res 50, D988–D995 (2022).

22. Lizio, M. et al. Gateways to the FANTOM5 promoter level mammalian expression atlas. Genome Biol 16, 22 (2015).

23. Rahman, F.A., Aziz, N. & Coverley, D. Differential detection of alternatively spliced variants of Ciz1 in normal and cancer cells using a custom exon-junction microarray. BMC Cancer 10, 482 (2010).

24. Rodermund, L. et al. Time-resolved structured illumination microscopy reveals key principles of Xist RNA spreading. Science 372 (2021).

25. Hall, L.L., Byron, M., Pageau, G. & Lawrence, J.B. AURKB-mediated effects on chromatin regulate binding versus release of XIST RNA to the inactive chromosome. J Cell Biol 186, 491–507 (2009).

26. Turvey, G.L. et al. Dominant CIZ1 fragments drive epigenetic instability and are expressed in early stage cancers. (BioRxiv).

27. Coleman, R.T. & Struhl, G. Causal role for inheritance of H3K27me3 in maintaining the OFF state of a Drosophila HOX gene. Science 356 (2017).

28. Jadhav, U. et al. Replicational Dilution of H3K27me3 in Mammalian Cells and the Role of Poised Promoters. Mol Cell 78, 141–151 e145 (2020).

29. Conway, E. et al. BAP1 enhances Polycomb repression by counteracting widespread H2AK119ub1 deposition and chromatin condensation. Mol Cell 81, 3526–3541 e3528 (2021).

30. Altun, M. et al. Activity-based chemical proteomics accelerates inhibitor development for deubiquitylating enzymes. Chem Biol 18, 1401–1412 (2011).

31. Lim, E. et al. Transcriptome analyses of mouse and human mammary cell subpopulations reveal multiple conserved genes and pathways. Breast Cancer Res 12, R21 (2010).

32. Thennavan, A. et al. Molecular analysis of TCGA breast cancer histologic types. Cell Genom 1 (2021).

33. Grimwood, J. et al. The DNA sequence and biology of human chromosome 19. Nature 428, 529–535 (2004).

34. Tate, J.G. et al. COSMIC: the Catalogue Of Somatic Mutations In Cancer. Nucleic Acids Res 47, D941–D947 (2019).

35. Meszaros, B. et al. PhaSePro: the database of proteins driving liquid-liquid phase separation. Nucleic Acids Res 48, D360–D367 (2020).

36. Quinodoz, S.A. et al. RNA promotes the formation of spatial compartments in the nucleus. Cell 184, 5775–5790 e5730 (2021).

37. You, J.S. & Jones, P.A. Cancer genetics and epigenetics: two sides of the same coin? Cancer Cell 22, 9–20 (2012).

38. Mack, S.C. et al. Epigenomic alterations define lethal CIMP-positive ependymomas of infancy. Nature 506, 445–450 (2014).

39. Dean, P.N. & Jett, J.H. Mathematical analysis of DNA distributions derived from flow microfluorometry. J Cell Biol 60, 523–527 (1974).

40. Livak, K.J. & Schmittgen, T.D. Analysis of relative gene expression data using real-time quantitative PCR and the 2(-Delta Delta C(T)) Method. Methods 25, 402–408 (2001).

41. Kim, D., Langmead, B. & Salzberg, S.L. HISAT: a fast spliced aligner with low memory requirements. Nat Methods 12, 357–360 (2015).

42. Pertea, M. et al. StringTie enables improved reconstruction of a transcriptome from RNA-seq reads. Nat Biotechnol 33, 290–295 (2015).

43. Subramanian, A. et al. Gene set enrichment analysis: a knowledge-based approach for interpreting genome-wide expression profiles. Proc Natl Acad Sci U S A 102, 15545–15550 (2005).

44. Xiao, Y. et al. A novel significance score for gene selection and ranking. Bioinformatics 30, 801–807 (2014).

45. Li, H. et al. The Sequence Alignment/Map format and SAMtools. Bioinformatics 25, 2078–2079 (2009).

46. Quinlan, A.R. & Hall, I.M. BEDTools: a flexible suite of utilities for comparing genomic features. Bioinformatics 26, 841–842 (2010).

47. Veiga, D.F.T. et al. A comprehensive long-read isoform analysis platform and sequencing resource for breast cancer. Sci Adv 8, eabg6711 (2022).

48. Bray, N.L., Pimentel, H., Melsted, P. & Pachter, L. Near-optimal probabilistic RNA-seq quantification. Nat Biotechnol 34, 525–527 (2016).

49. Soneson, C., Love, M.I. & Robinson, M.D. Differential analyses for RNA-seq: transcript-level estimates improve gene-level inferences. F1000Res 4, 1521 (2015).

50. Tang, Z. et al. GEPIA: a web server for cancer and normal gene expression profiling and interactive analyses. Nucleic Acids Res 45, W98–W102 (2017).

51. Kent, W.J. et al. The human genome browser at UCSC. Genome Res 12, 996–1006 (2002).

52. Bairoch, A. The Cellosaurus, a Cell-Line Knowledge Resource. J Biomol Tech 29, 25–38 (2018).

53. Fang, Z., Liu, X. & Peltz, G. GSEApy: a comprehensive package for performing gene set enrichment analysis in Python. Bioinformatics 39 (2023).

54. 54. Schindelin, J., et al. Fiji: an open-source platform for biological-image analysis. Nat Methods 9, 676–682 (2012).

55. Dahmcke, C.M., Buchmann-Moller, S., Jensen, N.A. & Mitchelmore, C. Altered splicing in exon 8 of the DNA replication factor CIZ1 affects subnuclear distribution and is associated with Alzheimer’s disease. Mol Cell Neurosci 38, 589–594 (2008).

56. Higgins, G. et al. Variant Ciz1 is a circulating biomarker for early-stage lung cancer. Proc Natl Acad Sci U S A 109, E3128–3135 (2012).

57. Rahman, F.A., Ainscough, J.F.-X., Copeland, N. & Coverley, D. Cancer-associated missplicing of exon 4 influences the subnuclear distribution of the DNA replication factor Ciz1. Human Mutation 28, 993–1004 (2007).

58. Swarts, D.R.A., Stewart, E.R., Higgins, G.S. & Coverley, D. CIZ1-F, an alternatively spliced variant of the DNA replication protein CIZ1 with distinct expression and localisation, is overrepresented in early stage common solid tumours. Cell Cycle, 1–16 (2018).

59. Plasari, G. et al. Nuclear factor I-C links platelet-derived growth factor and transforming growth factor beta1 signaling to skin wound healing progression. Mol Cell Biol 29, 6006–6017 (2009).

