## Supplemental material for "Epigenetic deprogramming driven by disruption of CIZ1-RNA nuclear assemblies in early-stage breast cancers"

**Supplemental Figure 1 (related to Fig.1). CIZ1 transcript in breast cancer cells**
A) Protein domain map aligning human (NP_001124488.1) and mouse (NP_082688.1) CIZ1. Numbers correspond to amino-acids encoded at exon boundaries. The domains highlighted are: Prion like domains 1 and 2 (PLD1 and PLD2, purple) at position 1-78 and 360-451 respectively (human) and position 1-67 and 361-399 respectively (mouse) [1], three zinc fingers (ZnF_C2H2 SM00355, ZF_C2H2 sd00020 and ZF_C2H2 sd00020, blue) at positions 593-617, 656-676 and 687-709 respectively (human) and 537-561, 600-620 and 631-653 respectively (mouse), an acidic domain (red) containing a concentrated area of aspartates and glutamates at position 741-761 (human) and 689-709 (mouse), and a matrin-3 homology domain (ZnF_U1 smart0045, yellow) at position 796-831 (human) and 746-770 (mouse). Boxes show % identity at the amino-acid level across these domains. Human and mouse CIZ1 are 65% identical at the protein level, with identity concentrated in the conserved domains (up to 96%).
B) CIZ1 exon-specific TPMs from four breast cancer (MCF7, BT-474, SK-BR-3 and MDA-MB-231) and one normal breast-tissue derived cell model (MCF10A), normalised to the first translated exon (exon 2), showing imbalanced domain expression, favouring the C-terminal anchor domain (AD). Below, exon map aligned with protein domains and location of epitopes detected throughout.
C) Frequency of reads aligning to human CIZ1 exon 10, demonstrating consistent coverage in the normal MCF10A line, and a transition in the cancer cell lines within exon 10, at the location of an alternative transcription start site.
D) *CIZ1* locus in *Homo sapiens* with corresponding exon numbers. Potential *CIZ1* alternative transcription start sites (TSSs) in exons 10 and 11 predicted in the FANTOM5 project [2] are indicated (red stars). The chromatin landscape in human mammary epithelial cells (HMEC), a cervical cancer cell line (HeLa) and a breast cancer cell line (MCF7) is shown below. Diagram generated using UCSC genome browser [3].

**Supplemental Figure 2. (related to Fig.2). CIZ1 domain expression in common solid tumours**

A) The location of amplicons detected by quantitative RT-PCR detection tools are shown in Fig.2A. These are four Taqman primer/probe sets; DT5 and DT7 which detect the 5' end of CIZ1 transcripts, and DT14 and DT16 which detect the 3' end. Dot plot shows comparison of outputs with the indicated pairs applied to 46 human tissue-derived cDNAs. Pearson's correlation coefficients show strong agreement between exons 5 and 7, and between 14 and 16, but poor agreement between exons 7 and 16, or 5 and 14, indicating that the 5' and 3' ends of CIZ1 are typically imbalanced at the transcript level.
B) Left, relative expression of exons 7 (red) and 16 (blue), normalized to the average of 3 unmatched control samples for each of six common solid tumour types in multi-tissue cDNA array CSRT101 aggregated by disease stage (0-IV, where 0 represents histologically normal tissue). Middle, individual sample data plus the average of the controls calibrated to 1 (Av.C). Right, plots show the ratio of exon 7 to exon 16 ordered by stage, with trendline (polynomial 2) indicative of a shift in ratio at stage IV. Individual sample information for all arrays in given in Supplemental Dataset 2.
C) Total CIZ1 TPM derived from the indicated number of cancer (C) and normal (N) tissues in TCGA, compared using GEPIA for the indicated disease types. No significant difference is detected (where $log_2FC$ was >1, and p-value < 0.05), when comparing all amalgamated transcripts that map to the CIZ1 gene (unresolved by exon).

**Supplemental Figure 3 (related to Fig.3). Cell cycle analysis and anchor domain mutagenesis**

A) Example image showing accumulation of GFP-C275 (green) at the site of endogenous CIZ1-Xi assemblies (red). DNA is blue, bar is 1µm. Right, frequency of cells with C275 enrichment at sites of endogenous CIZ1-Xi assemblies in D3T3 cells at 24 hours after transfection. Across serial dilution cultures the frequency increases with cell density and contact inhibition, illustrating requirement for growing cultures.
B) CIZ1-Xi assembly frequency 1-5 hours after release from cell cycle arrest in S phase (thymidine) or M phase (nocodazole). N is 2-4 as indicated, number of nuclei inspected at each time point is given (n). Comparisons between 1 and 5 hours t-test, where p=0.45 for S phase and $p=1.15x10^{-6}$ for M-G1 phase.
C) Map of C181 deletion constructs, showing excluded sequences in single letter code. These exclude the MH3 domain (conserved domain ZnF_U1 smart00451), or the fully human/mouse conserved sequence downstream of the MH3 domain (ΔNALTALF), or the murine equivalent of the 8 amino acids previously implicated in lung cancer (CIZ1B) [4], or the terminal 37 amino acids (Δ37). We also evaluated a fragment encompassing the MH3 domain but lacking sequences up and downstream (I122) [5]. Numbers indicate amino-acid at boundaries relative to murine full-length CIZ1. AcD, acidic domain (red), MH3, matrin 3 homology domain (yellow).
D) Field views of D3T3 cells expressing GFP-tagged C181-derived deletion mutants, without (total) and with (detergent-resistant) pre-fixation wash with 0.05% Triton X-100. The percentages show the proportion of transfected cells in each population with nuclear C181 or derivative, revealing the degree of sensitivity to extraction. Bar is 5µm.
E) Effect of fragments on the frequency of endogenous CIZ1-Xi assemblies in D3T3 cells. Compared to C181, only ΔMH3 was perturbed in its ability to disperse endogenous CIZ1 (p=0.011). All other deletion mutants retained similar DNF capability to C181 (ΔNALTALF p=0.96, Δ37 p=0.64, CIZ1B p=0.99, I122 p=1). N shows replicate analyses with total nuclei inspected in parentheses. Comparisons are by one-way ANOVA. Error bars show SEM.
F) Left, box and whisker plot showing normalized endogenous CIZ1-RD fluorescence intensity per nucleus in female WT PEFs, either untransfected (UT) or with C181, or derived deletion mutant ΔMH3, showing reduced potency of ΔMH3 compared to C181 (p=0.002, t-test). Right, mean intensity measures ordered low to high for endogenous CIZ1 in UT, C181 and ΔMH3 expressing cells. Mean values are also shown. Below, example images of cells stained for endogenous CIZ1 (red), with and without ectopic GFP-C181 or GFP-ΔMH3. Bar is 5µm. Inset shows surviving Xi assemblies in grey scale.

**Supplemental Figure 4 (related to Fig.4). Effect of CIZ1 assembly dispersal on H3K27me3 and H2AK119ub1**

A) Graphs show the frequency of endogenous CIZ1-Xi assemblies in a cycling population of female D3T3 cells, comparing transfected and untransfected cells in the same population. Endogenous CIZ1 assemblies are detected via CIZ1-RD and classified into three categories; present, absent or intermediate. Middle and lower graphs show the frequency of repressive histone marks in cells that are, or are not transfected with GFP-C181. N is replicate analyses, with nuclei scored in parentheses. Comparisons are by t-test. For endogenous CIZ1 in UT and C181 cells p=0.00023, for H3K27me3 p=0.60, for H2AK119ub1 p=0.0073. Error bars show SEM.
B) As in A, except that all data is derived from analysis of female primary embryonic fibroblasts (PEFs) at passage 2-3. For endogenous CIZ1 in UT and C181 cells p=0.00033, for H3K27me3 p=0.79, for H2AK119ub1 p=0.016, performed on present (type 1) categories. Also shown, the effect of 5µM PR619 on H2AK119ub1 loss, where p=0.0099 for the no CIZ1 category (type 3). Error bars show SEM.
C) Example images of endogenous CIZ1 and histone marks (red) in untransfected (UT) and C181 transfected (green) WT PEF populations. Bar is 10µm.
D) Possible mechanisms by which CIZ1 assemblies might influence H2AK119ub1 dynamics on Xi chromatin. Recruitment model: CIZ1-*Xist* assemblies contribute to recruitment or activation of PRC1, supporting H2AK119ub1 deposition. Shield model: Multiple CIZ1 dimers and RNAs coalesce to form a molecular shield at the Xi which blocks access to deubiquitinating enzymes, supporting H2AK119ub1 preservation.
E) Frequency of cells with H2AK119ub1 at the Xi in the vector only population, and cells transduced with C181, without and with the DUB inhibitor PR619 (5 µM). N is replicate analyses, with total nuclei inspected in parentheses. Comparisons are by unpaired t-test. Error bars show SEM. Below, example images taken under standardised conditions showing H2AK119ub1 in red in C181 transduced WT primary embryonic fibroblasts.

F) Restoration of H2AK119ub1 enrichment at Xi in CIZ1 null primary embryonic fibroblasts by PR619. In untreated cells approximately 10% of cells have H2AK119ub1 enriched Xi's, which increased to approximately 35% within 24 hours of treatment, while H3K27me3 remains unchanged. Error bars show SEM. Right, example images of H2AK119ub1 in CIZ1 null primary embryonic fibroblasts.

**Supplemental Figure 5 (related to Fig.4). Effect of CIZ1 assembly dispersal on gene expression**

A) Principle component analysis showing clustering of triplicate PEF-derived transcriptomes for each condition.
B) Scatter plot showing mean transcripts per million reads (TPM), coloured blue (down, DN) and red (UP) for genes meeting the false-detection rate corrected q values of <0.05 and log2 FC 1.
C) Example gene set enrichment analysis performed on pre-ranked transcription units derived by comparing expression in C181 transduced cells to the empty vector control, showing highly significant enrichment in two breast stem and cancer-related gene sets, and two controlled by TGfb.
D) Chromosomal locations of C181-driven UP regulated genes (orange) and DN regulated genes (grey, FDR q<0.05, log2FC1), with centromeres shown in black. Circled are regions of synteny to the human cluster regions highlighted in Fig.6. Human cluster at 1q21.3 (148-158,000,000) is syntenic with murine chromosome 3 (86,903,019-97,986,449), human cluster at 6p21.31 (250-350,000,000) is syntenic with murine chromosome 17 (17: 27,135,758-31,159,854), human cluster at 9q34 (128-138,000,000) is syntenic with murine chromosome 2 (24,493,799-32,150,031), human cluster at 10q22.2 (70-80,000,000) is syntenic with murine chromosome 14 (20,344,703-25,806,867), human cluster at 11q13.1 (60-70,000,000) is syntenic with murine chromosome 19 (3,309,831-13,840,444), human cluster at 16p13.3 (0-10,000000) is syntenic with murine chromosome 17 (23,765,442-26,506,126).

**Supplemental Figure 6 (related to Fig.5). Segmentation of TCGA transcriptomes.**

A) TCGA breast cancers by histological subtype [6], showing representation across the DNF index as variance from the full breast cancer TCGA cohort profile. The number of samples in each subtype is denoted by n. Similarity to cohort profile was evaluated by Chi squared test, where P<0.05 is considered significant. Dotted line indicates lack of variance.
B) As in A, but after classification by receptor status [6].
C) Variance between within receptor positive and receptor negative groups.
D) As in A, but after classification by tumour stage.
E) Top 10 GSEA curated gene sets (M2 CGP) returned by DN genes, including 3 related to response to UV, and two related to apoptosis.
D) Top 20 GSEA curated gene sets (M2 CGP) returned by UP genes, including 6 breast cancer related sets, one describing mammary stem cell phenotype, and one describing genes normally suppressed by PRC2 catalytic subunit EZH2.
E) Top 10 GSEA cell type signatures (C8) returned by UP genes, including primarily fetal cell types but no normal breast tissue signatures.

**Supplemental Figure 7 (related to Fig.6).**

A) Upper, differentially expressed lncRNAs on chromosome 1 (green) returned by comparison of TCGA breast tumours with DNF index A (elevated AD), compared to C (balanced RD and AD). Unaffected genes are shown in grey. Middle, as above for protein coding genes (yellow). Lower, chromatin accessibility revealed by ATACseq in 8 group A tumours compared to 15 group C tumours, with non-significant intervals in grey and differentially accessible intervals in blue. ATACseq peaks indicate are evident across all chromosomes, and within cluster regions are exclusively UP.
Cluster region is marked with a box (10Mb).
B) As in A, for chromosome 6.
C) As in A, for chromosome 9.
D) As in A, for chromosome 10.
E) As in A, for chromosome 11.
F) As in A, for chromosome 16.
G) Table showing summary of gene densities across the indicated 10Mbp domains, relative to overall density on each analysed chromosome. UP clusters encode more genes (coding and non-coding) than the chromosomal averages, are similarly represented in down-regulated protein coding genes but over-represented in up-regulated protein coding genes. For lncRNAs, both UP and DN genes are enriched in excess of the gene density.

1. Sofi, S. *et al.* Prion-like domains drive CIZ1 assembly formation at the inactive X chromosome. *J Cell Biol* **221** (2022).
2. Lizio, M. *et al.* Gateways to the FANTOM5 promoter level mammalian expression atlas. *Genome Biol* **16**, 22 (2015).
3. Kent, W.J. *et al.* The human genome browser at UCSC. *Genome Res* **12**, 996-1006 (2002).
4. Higgins, G. *et al.* Variant Ciz1 is a circulating biomarker for early-stage lung cancer. *Proc Natl Acad Sci U S A* **109**, E3128-3135 (2012).
5. Ainscough, J.F. *et al.* C-terminal domains deliver the DNA replication factor Ciz1 to the nuclear matrix. *Journal of Cell Science* **120**, 115-124 (2007).
6. Thennavan, A. *et al.* Molecular analysis of TCGA breast cancer histologic types. *Cell Genom* **1** (2021).

### SFig.1

A. Conserved sequence elements with validated or predicted functions

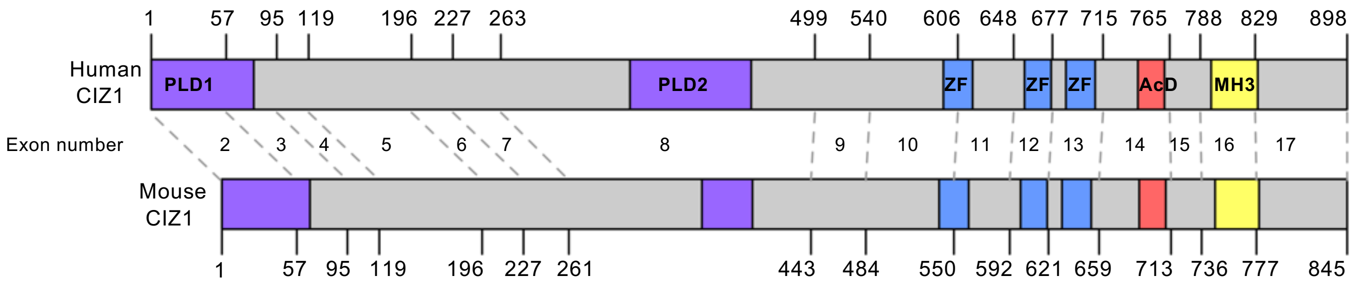


| Domain | PLD1 | PLD2 | ZF1 | ZF2 | ZF3 | AcD | MH3 |
|---|---|---|---|---|---|---|---|
| Percentage identity | 78 | 71 | 92 | 86 | 96 | 67 | 79 |

B. Relative exon abundance in breast-derived cell lines

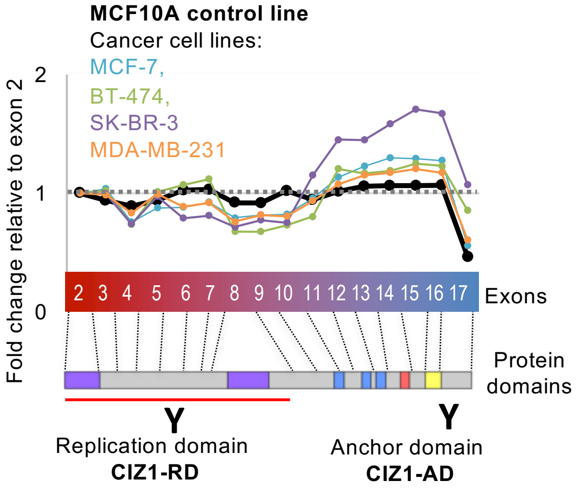


C. A transition within human CIZ1 exon 10

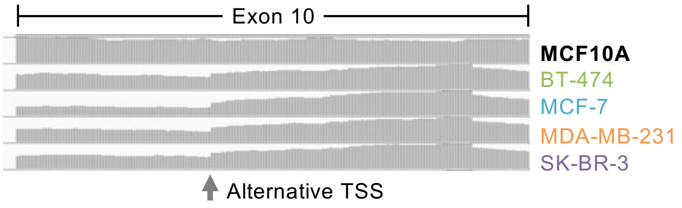


D. Epigenetic landscape of the human *CIZ1* locus in normal and cancer cells

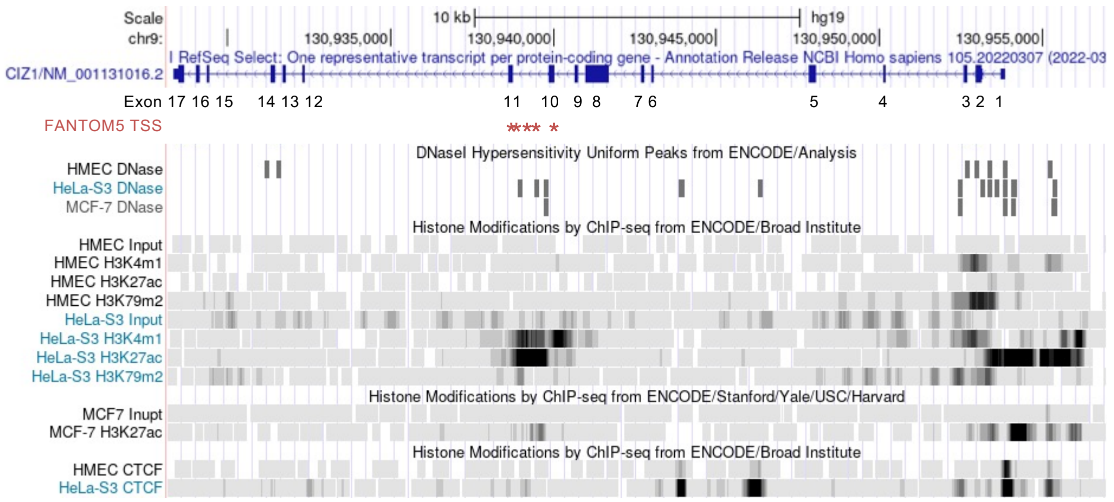

### SFig.2

A. Comparison of RD and AD amplicon detection

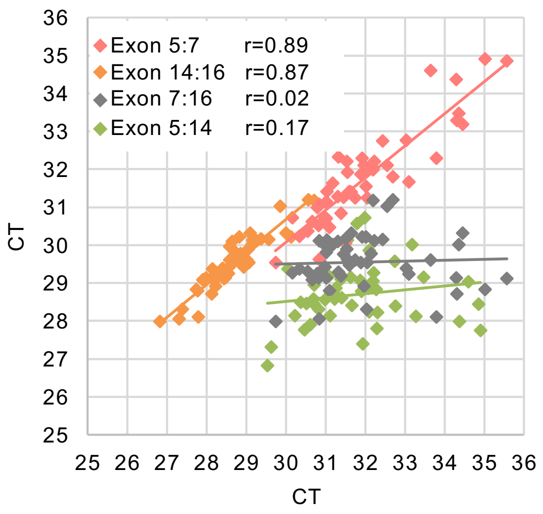


C. Total CIZ1 TPM in TCGA tumour and TCGA normal tissues

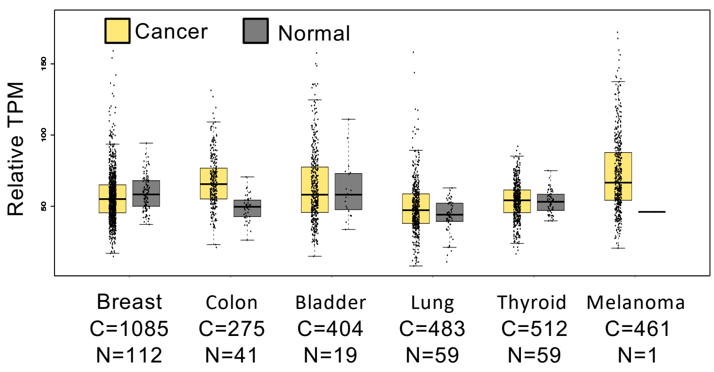


B. RD and AD amplicon expression aggregated by tumour stage

RD and AD (levels in Individual patients)

Ratio: RQ exon 7:16

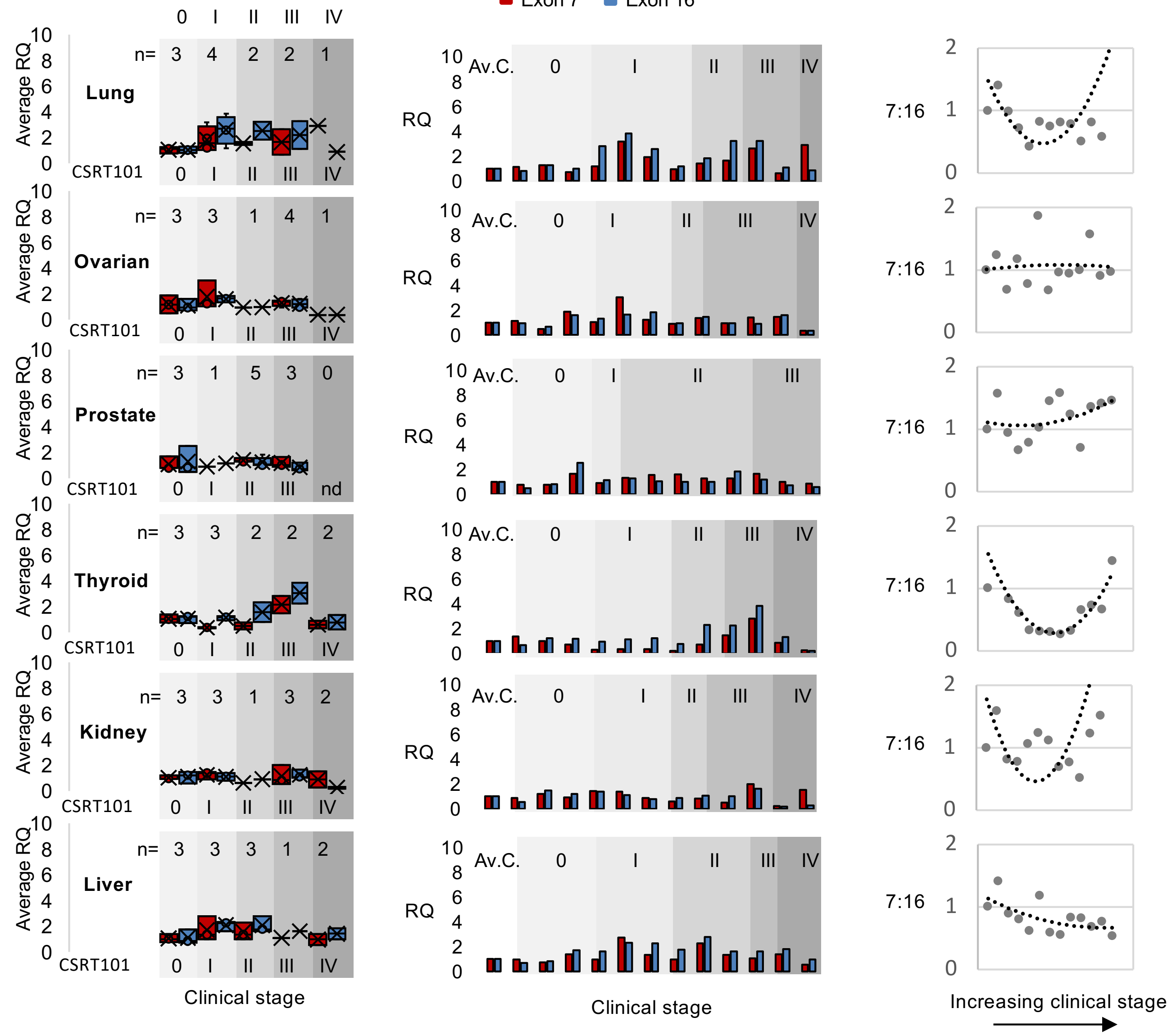

### SFig.3

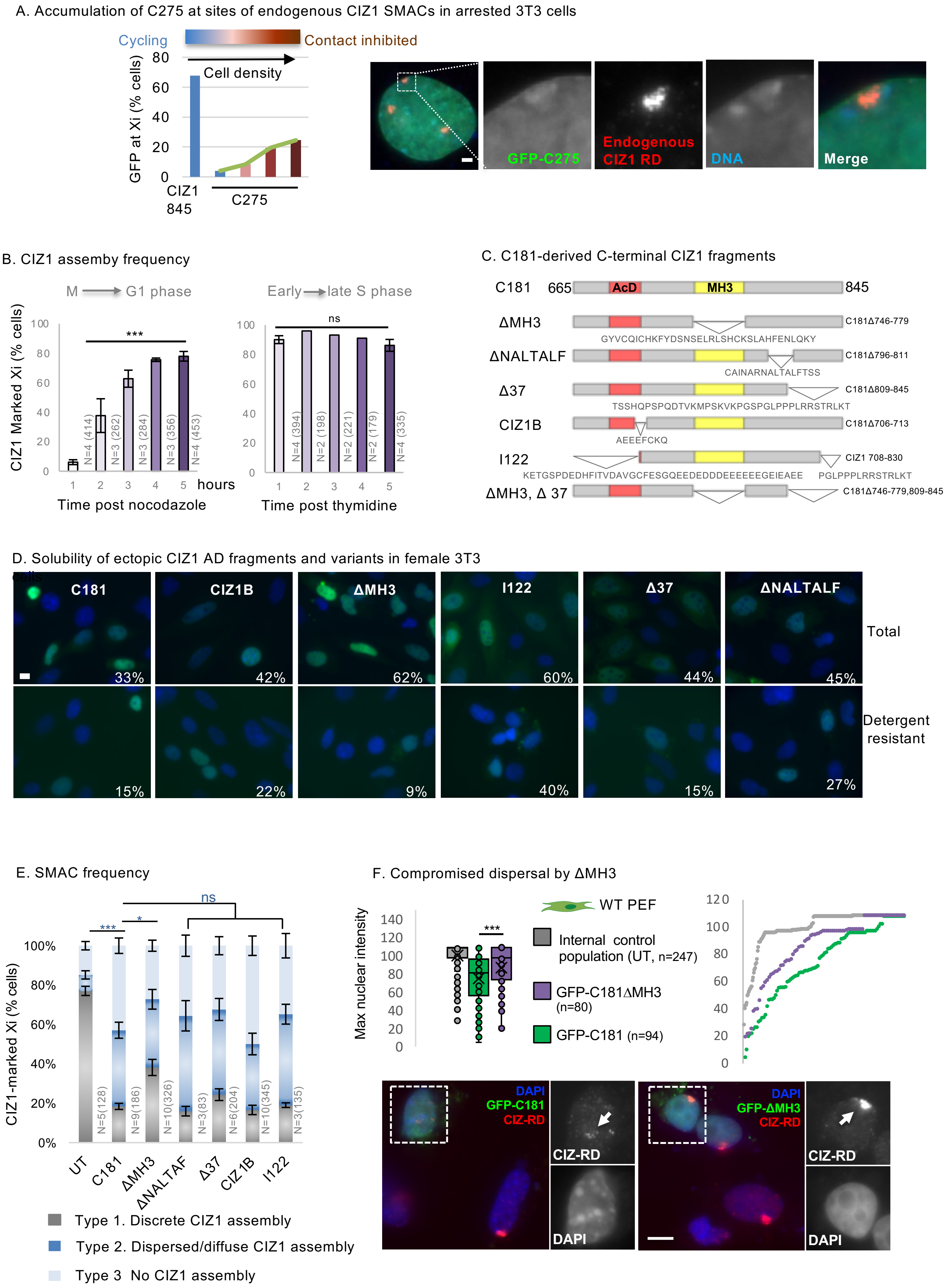
A. Accumulation of C275 at sites of endogenous CIZ1 SMACs in arrested 3T3 cells
Cycling
Contact inhibited
Cell density
GFP at Xi (% cells)
80
60
40
20
0
CIZ1 845
C275
GFP-C275
Endogenous CIZ1 RD
DNA
Merge
B. CIZ1 assemby frequency
M → G1 phase
Early → late S phase
CIZ1 Marked Xi (% cells)
100
80
60
40
20
0
***
ns
N=4 (414)
N=3 (262)
N=3 (284)
N=3 (356)
N=4 (453)
N=4 (394)
N=2 (198)
N=2 (221)
N=2 (179)
N=4 (335)
1 2 3 4 5 hours
Time post nocodazole
Time post thymidine
C. C181-derived C-terminal CIZ1 fragments
C181 665 AcD MH3 845
ΔMH3 C181Δ746-779
GYVCQICHKFYDSNSELRLSHCKSLAHFENLQKY
ΔNALTALF C181Δ796-811
CAINARNALTALFTSS
Δ37 C181Δ809-845
TSSHQPSPQDTVKMPSKVKPGSPGLPPPLRRSTRLKT
CIZ1B C181Δ706-713
AEEEFCKQ
I122 CIZ1 708-830
KETGSPDEDHFITVDAVGCFESGQEEDEDDDEEEEEEGEIEAEE PGLPPPLRRSTRLKT
ΔMH3, Δ 37 C181Δ746-779,809-845
D. Solubility of ectopic CIZ1 AD fragments and variants in female 3T3 cells
C181
CIZ1B
ΔMH3
I122
Δ37
ΔNALTALF
33%
42%
62%
60%
44%
45%
Total
15%
22%
9%
40%
15%
27%
Detergent resistant
E. SMAC frequency
ns
***
*
CIZ1-marked Xi (% cells)
100%
80%
60%
40%
20%
0%
N=5(128)
N=9(186)
N=10(326)
N=3(83)
N=6(204)
N=10(345)
N=3(135)
UT
C181
ΔMH3
ΔNALTAF
Δ37
CIZ1B
I122
Type 1. Discrete CIZ1 assembly
Type 2. Dispersed/diffuse CIZ1 assembly
Type 3 No CIZ1 assembly
F. Compromised dispersal by ΔMH3
WT PEF
Max nuclear intensity
140
120
100
80
60
40
20
0
***
Internal control population (UT, n=247)
GFP-C181ΔMH3 (n=80)
GFP-C181 (n=94)
120
100
80
60
40
20
0
DAPI
GFP-C181
CIZ-RD
CIZ-RD
DAPI
DAPI
GFP-ΔMH3
CIZ-RD
CIZ-RD
DAPI

### SFig.4

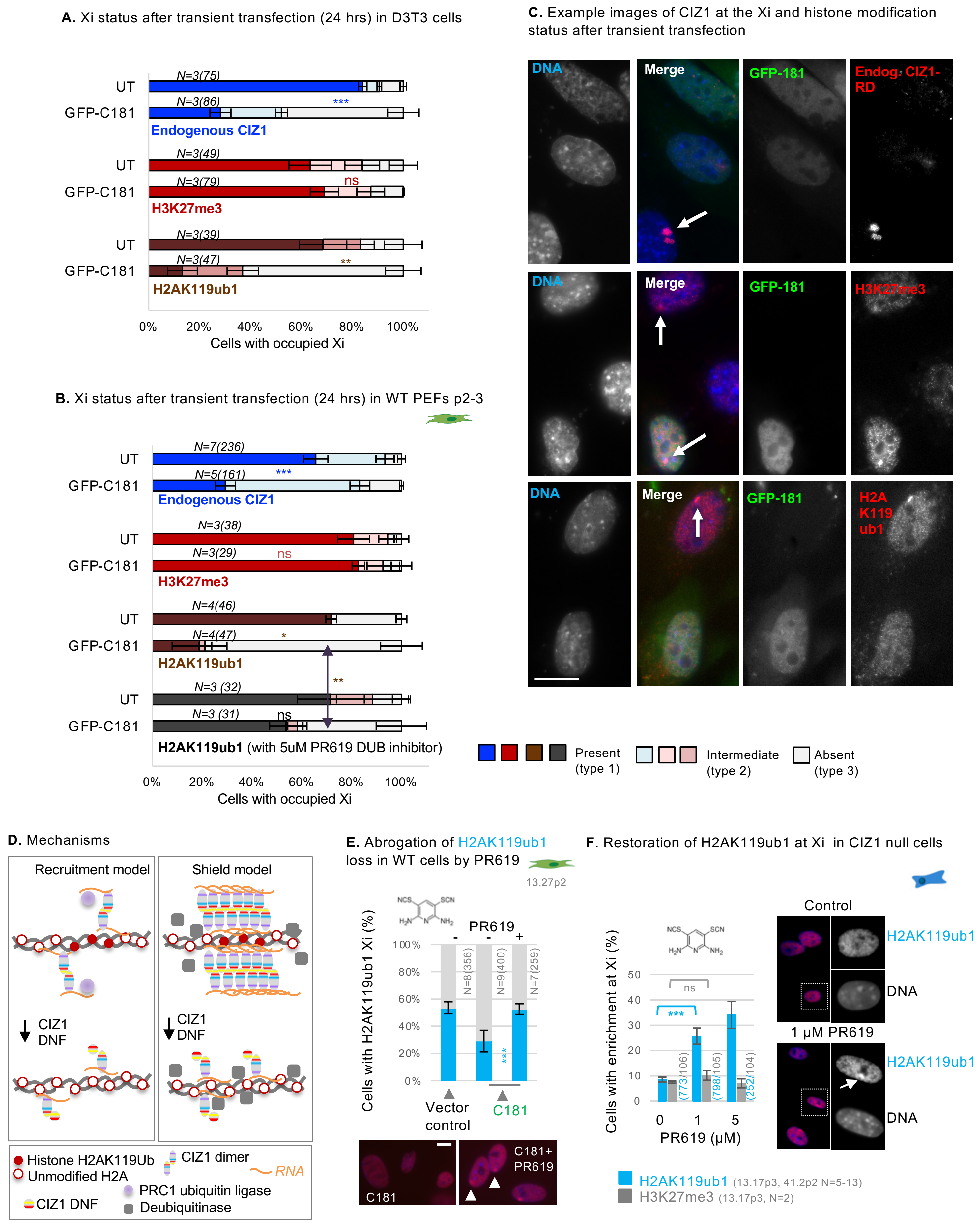

A. Xi status after transient transfection (24 hrs) in D3T3 cells
UT
GFP-C181
N=3(75)
N=3(86)
***
Endogenous CIZ1
N=3(49)
N=3(79)
ns
H3K27me3
N=3(39)
N=3(47)
**
H2AK119ub1
0%
20%
40%
60%
80%
100%
Cells with occupied Xi
B. Xi status after transient transfection (24 hrs) in WT PEFs p2-3
N=7(236)
N=5(161)
***
Endogenous CIZ1
N=3(38)
N=3(29)
ns
H3K27me3
N=4(46)
N=4(47)
*
H2AK119ub1
**
N=3 (32)
N=3 (31)
ns
H2AK119ub1 (with 5uM PR619 DUB inhibitor)
Cells with occupied Xi
C. Example images of CIZ1 at the Xi and histone modification status after transient transfection
DNA
Merge
GFP-181
Endog. CIZ1-RD
H3K27me3
H2A K119 ub1
Present (type 1)
Intermediate (type 2)
Absent (type 3)
D. Mechanisms
Recruitment model
Shield model
CIZ1 DNF
Histone H2AK119Ub
Unmodified H2A
CIZ1 DNF
CIZ1 dimer
RNA
PRC1 ubiquitin ligase
Deubiquitinase
E. Abrogation of H2AK119ub1 loss in WT cells by PR619
13.27p2
PR619
- - +
100%
80%
60%
40%
20%
0%
N=8(356)
N=9(400)
N=7(259)
***
Cells with H2AK119ub1 Xi (%)
Vector control
C181
C181
C181+ PR619
F. Restoration of H2AK119ub1 at Xi in CIZ1 null cells
Cells with enrichment at Xi (%)
50
40
30
20
10
0
ns
***
(773/106)
(798/105)
(252/104)
0 1 5
PR619 (μM)
Control
H2AK119ub1
DNA
1 μM PR619
H2AK119ub1
DNA
H2AK119ub1 (13.17p3, 41.2p2 N=5-13)
H3K27me3 (13.17p3, N=2)

### SFig.5

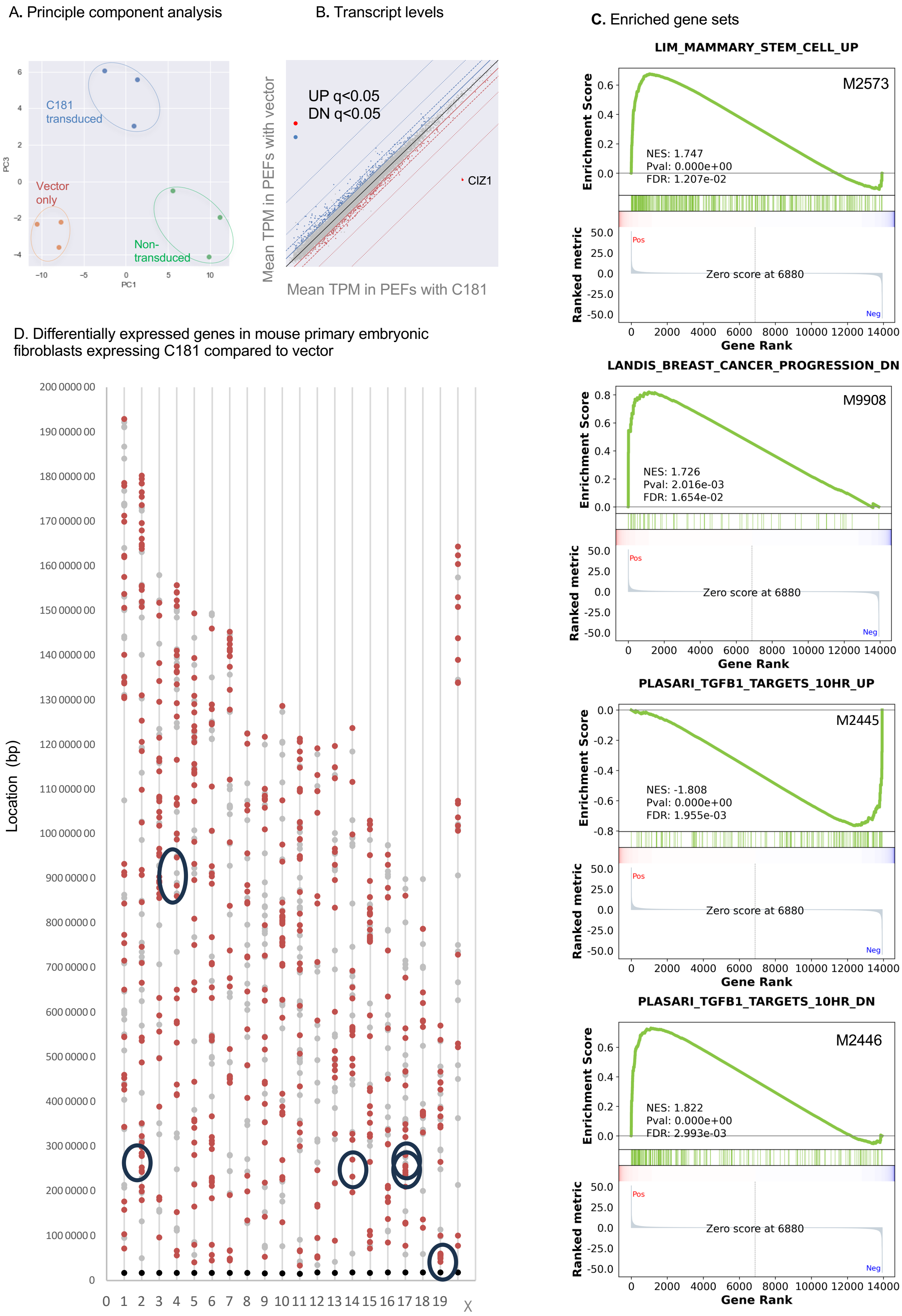

A. Principle component analysis
C181
transduced
Vector
only
Non-
transduced
PC1
PC3
B. Transcript levels
UP q<0.05
DN q<0.05
CIZ1
Mean TPM in PEFs with vector
Mean TPM in PEFs with C181
C. Enriched gene sets
LIM_MAMMARY_STEM_CELL_UP
M2573
NES: 1.747
Pval: 0.000e+00
FDR: 1.207e-02
LANDIS_BREAST_CANCER_PROGRESSION_DN
M9908
NES: 1.726
Pval: 2.016e-03
FDR: 1.654e-02
PLASARI_TGFB1_TARGETS_10HR_UP
M2445
NES: -1.808
Pval: 0.000e+00
FDR: 1.955e-03
PLASARI_TGFB1_TARGETS_10HR_DN
M2446
NES: 1.822
Pval: 0.000e+00
FDR: 2.993e-03
Enrichment Score
Ranked metric
Pos
Neg
Zero score at 6880
Gene Rank
D. Differentially expressed genes in mouse primary embryonic fibroblasts expressing C181 compared to vector
Location (bp)
0 1 2 3 4 5 6 7 8 9 10 11 12 13 14 15 16 17 18 19 X

### SFig.6

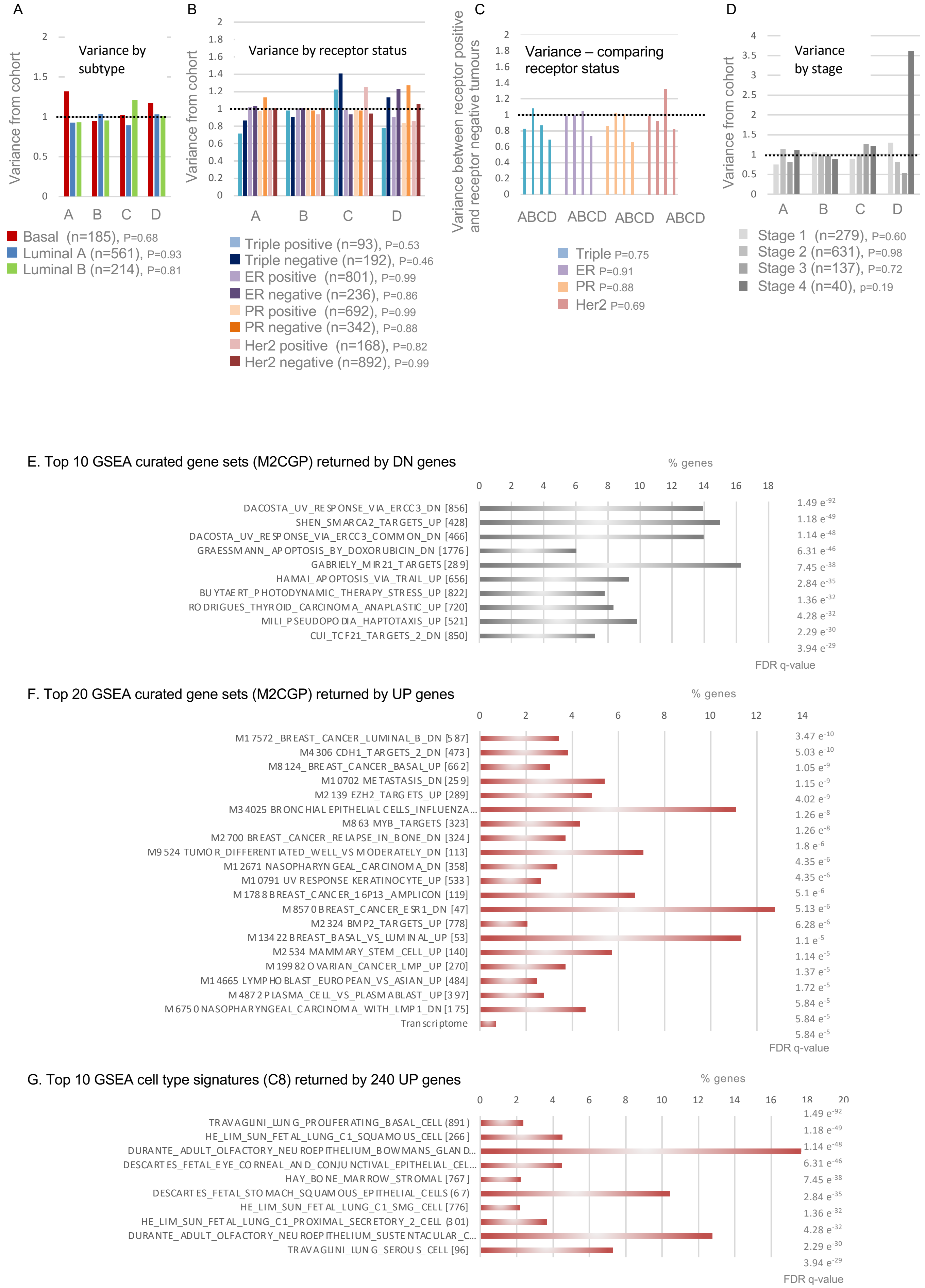

A
Variance by subtype
Variance from cohort
2
1.5
1
0.5
0
A B C D
Basal (n=185), P=0.68
Luminal A (n=561), P=0.93
Luminal B (n=214), P=0.81
B
Variance by receptor status
Variance from cohort
2
1.8
1.6
1.4
1.2
1
0.8
0.6
0.4
0.2
0
A B C D
Triple positive (n=93), P=0.53
Triple negative (n=192), P=0.46
ER positive (n=801), P=0.99
ER negative (n=236), P=0.86
PR positive (n=692), P=0.99
PR negative (n=342), P=0.88
Her2 positive (n=168), P=0.82
Her2 negative (n=892), P=0.99
C
Variance – comparing receptor status
Variance between receptor positive and receptor negative tumours
2
1.8
1.6
1.4
1.2
1
0.8
0.6
0.4
0.2
0
ABCD ABCD ABCD ABCD
Triple P=0.75
ER P=0.91
PR P=0.88
Her2 P=0.69
D
Variance by stage
Variance from cohort
4
3.5
3
2.5
2
1.5
1
0.5
0
A B C D
Stage 1 (n=279), P=0.60
Stage 2 (n=631), P=0.98
Stage 3 (n=137), P=0.72
Stage 4 (n=40), p=0.19
E. Top 10 GSEA curated gene sets (M2CGP) returned by DN genes
% genes
0 2 4 6 8 10 12 14 16 18
DACOSTA_UV_RESPONSE_VIA_ERCC3_DN [856]
SHEN_SMARCA2_TARGETS_UP [428]
DACOSTA_UV_RESPONSE_VIA_ERCC3_COMMON_DN [466]
GRAESSMANN_APOPTOSIS_BY_DOXORUBICIN_DN [1776]
GABRIELY_MIR21_TARGETS [289]
HAMAI_APOPTOSIS_VIA_TRAIL_UP [656]
BUYTAERT_PHOTODYNAMIC_THERAPY_STRESS_UP [822]
RODRIGUES_THYROID_CARCINOMA_ANAPLASTIC_UP [720]
MILI_PSEUDOPODIA_HAPTOTAXIS_UP [521]
CUI_TCF21_TARGETS_2_DN [850]
1.49 e-92
1.18 e-49
1.14 e-48
6.31 e-46
7.45 e-38
2.84 e-35
1.36 e-32
4.28 e-32
2.29 e-30
3.94 e-29
FDR q-value
F. Top 20 GSEA curated gene sets (M2CGP) returned by UP genes
% genes
0 2 4 6 8 10 12 14
M17572_BREAST_CANCER_LUMINAL_B_DN [587]
M4306 CDH1_TARGETS_2_DN [473]
M8124_BREAST_CANCER_BASAL_UP [662]
M10702 METASTASIS_DN [259]
M2139 EZH2_TARGETS_UP [289]
M34025 BRONCHIAL EPITHELIAL CELLS_INFLUENZA...
M863 MYB_TARGETS [323]
M2700 BREAST_CANCER_RELAPSE_IN_BONE_DN [324]
M9524 TUMOR_DIFFERENTIATED_WELL_VS MODERATELY_DN [113]
M12671 NASOPHARYNGEAL_CARCINOMA_DN [358]
M10791 UV RESPONSE KERATINOCYTE_UP [533]
M1788 BREAST_CANCER_16P13_AMPLICON [119]
M8570 BREAST_CANCER_ESR1_DN [47]
M2324 BMP2_TARGETS_UP [778]
M13422 BREAST_BASAL_VS_LUMINAL_UP [53]
M2534 MAMMARY_STEM_CELL_UP [140]
M19982 OVARIAN_CANCER_LMP_UP [270]
M14665 LYMPHOBLAST_EUROPEAN_VS_ASIAN_UP [484]
M4872 PLASMA_CELL_VS_PLASMABLAST_UP [397]
M6750 NASOPHARYNGEAL_CARCINOMA_WITH_LMP1_DN [175]
Transcriptome
3.47 e-10
5.03 e-10
1.05 e-9
1.15 e-9
4.02 e-9
1.26 e-8
1.26 e-8
1.8 e-6
4.35 e-6
4.35 e-6
5.1 e-6
5.13 e-6
6.28 e-6
1.1 e-5
1.14 e-5
1.37 e-5
1.72 e-5
5.84 e-5
5.84 e-5
5.84 e-5
FDR q-value
G. Top 10 GSEA cell type signatures (C8) returned by 240 UP genes
% genes
0 2 4 6 8 10 12 14 16 18 20
TRAVAGLINI_LUNG_PROLIFERATING_BASAL_CELL (891)
HE_LIM_SUN_FETAL_LUNG_C1_SQUAMOUS_CELL [266]
DURANTE_ADULT_OLFACTORY_NEUROEPITHELIUM_BOWMANS_GLAND...
DESCARTES_FETAL_EYE_CORNEAL_AND_CONJUNCTIVAL_EPITHELIAL_CEL...
HAY_BONE_MARROW_STROMAL [767]
DESCARTES_FETAL_STOMACH_SQUAMOUS_EPITHELIAL_CELLS (67)
HE_LIM_SUN_FETAL_LUNG_C1_SMG_CELL [776]
HE_LIM_SUN_FETAL_LUNG_C1_PROXIMAL_SECRETORY_2_CELL (301)
DURANTE_ADULT_OLFACTORY_NEUROEPITHELIUM_SUSTENTACULAR_C...
TRAVAGLINI_LUNG_SEROUS_CELL [96]
1.49 e-92
1.18 e-49
1.14 e-48
6.31 e-46
7.45 e-38
2.84 e-35
1.36 e-32
4.28 e-32
2.29 e-30
3.94 e-29
FDR q-value

### SFig.7

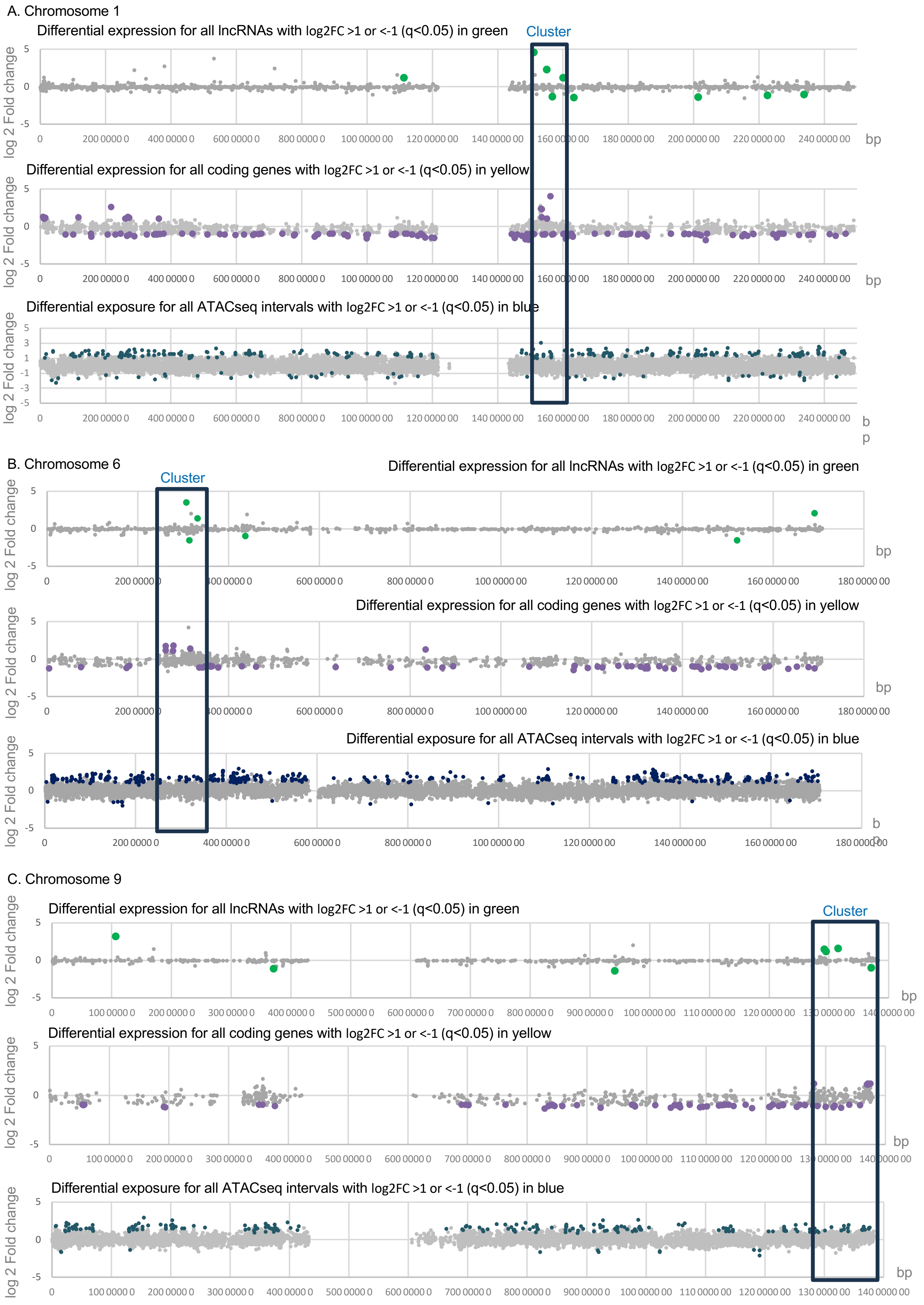

A. Chromosome 1
Differential expression for all lncRNAs with log2FC >1 or <-1 (q<0.05) in green
Cluster
log 2 Fold change
bp
Differential expression for all coding genes with log2FC >1 or <-1 (q<0.05) in yellow
Differential exposure for all ATACseq intervals with log2FC >1 or <-1 (q<0.05) in blue
0
200 0000 0
400 0000 0
600 0000 0
800 0000 0
100 0000 00
120 0000 00
140 0000 00
160 0000 00
180 0000 00
200 0000 00
220 0000 00
240 0000 00
B. Chromosome 6
Differential expression for all lncRNAs with log2FC >1 or <-1 (q<0.05) in green
Cluster
Differential expression for all coding genes with log2FC >1 or <-1 (q<0.05) in yellow
Differential exposure for all ATACseq intervals with log2FC >1 or <-1 (q<0.05) in blue
C. Chromosome 9
Differential expression for all lncRNAs with log2FC >1 or <-1 (q<0.05) in green
Cluster
Differential expression for all coding genes with log2FC >1 or <-1 (q<0.05) in yellow
Differential exposure for all ATACseq intervals with log2FC >1 or <-1 (q<0.05) in blue

### SFig.7

D. Chromosome 10

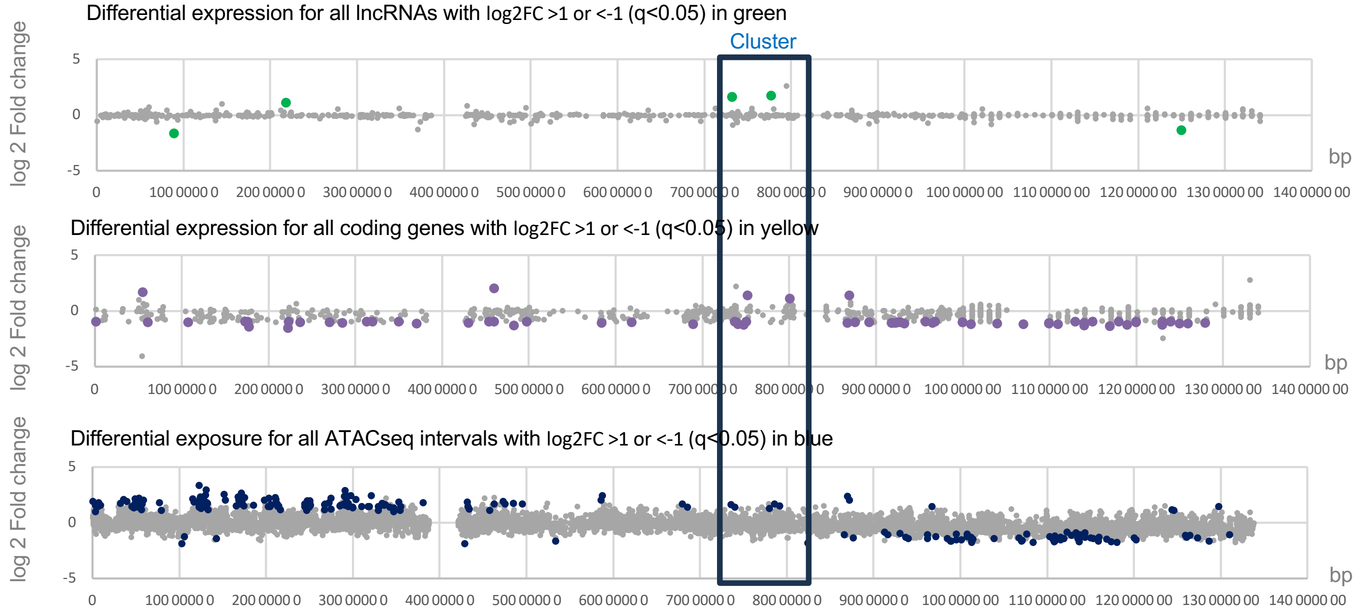


E. Chromosome 11

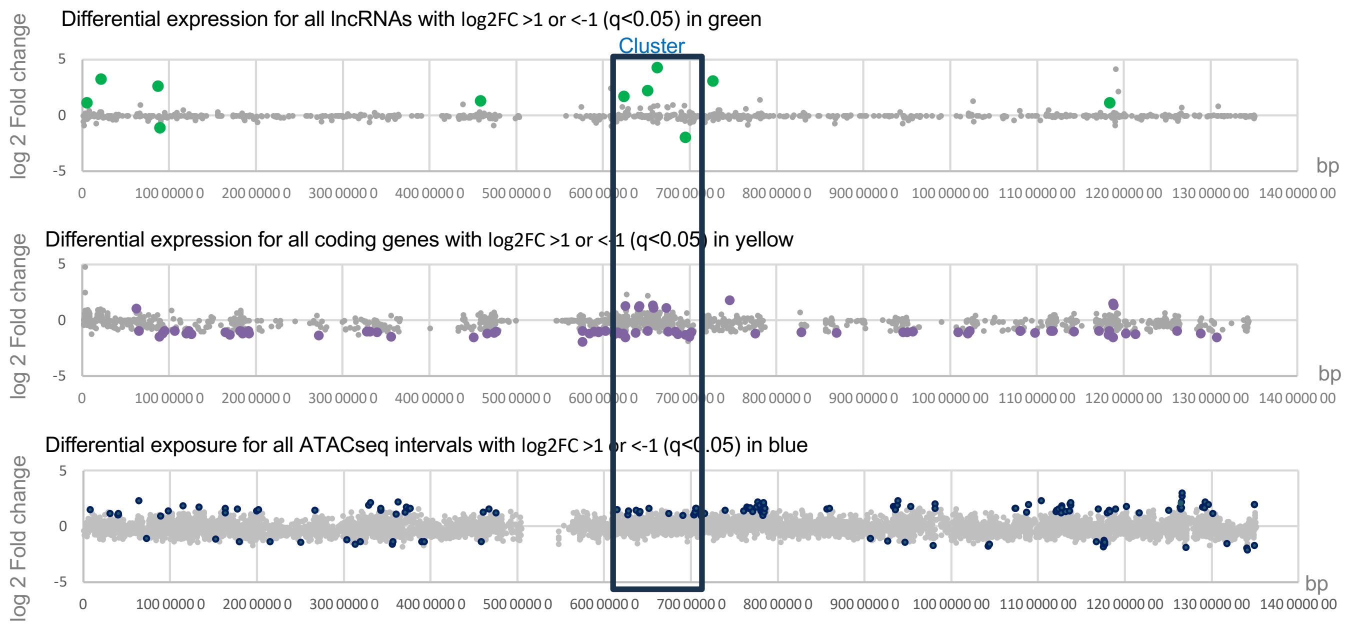


F. Chromosome 16

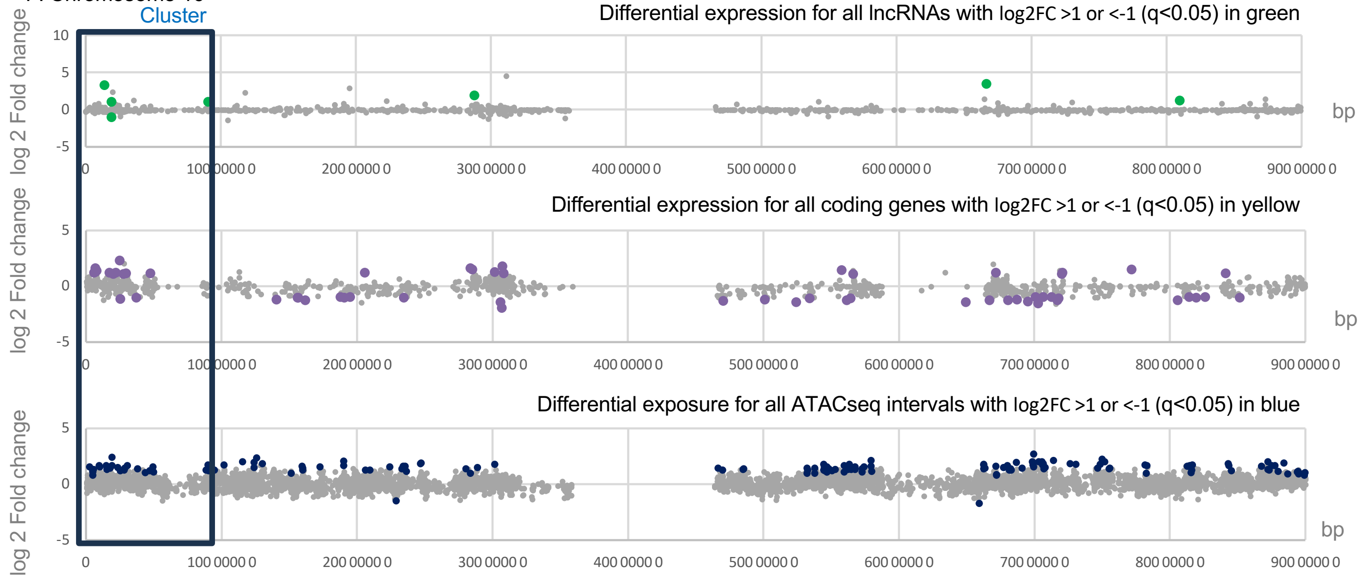

### SFig.7

G. Summary of gene densities across the indicated 10Mbp domain, relative to overall density on each analysed chromosome.

| | **Average spacing(Kb)** | | | **Average spacing(Kb)** | | | **Average spacing(Kb)** | |
|---|---|---|---|---|---|---|---|---|
| All genes | Protein coding | lncRNAs | UP DEGs | Protein coding | lncRNAs | DN DEGs | Protein coding | lncRNAs |
| Chr1 | 120.7 | 174.4 | Chr1 | 15453.1 | 61812.4 | Chr1 | 1873.1 | 49449.9 |
| Cluster 150-160 | 35.2 | 116.3 | Cluster 150-160 | 2000.0 | 3333.3 | Cluster 150-160 | 1428.6 | 10000.0 |
| fold enrichment | 3.4 | 1.5 | fold enrichment | 7.7 | 18.5 | fold enrichment | 1.3 | 4.9 |

| All genes | Protein coding | lncRNAs | UP DEGs | Protein coding | lncRNAs | DN DEGs | Protein coding | lncRNAs |
|---|---|---|---|---|---|---|---|---|
| Chr6 | 163.2 | 205.4 | Chr6 | 28483.3 | 56966.7 | Chr6 | 3224.5 | 56966.7 |
| Cluster 250-350 | 34.2 | 96.2 | Cluster 250-350 | 2000.0 | 5000.0 | Cluster 250-350 | 2000.0 | 10000.0 |
| fold enrichment | 4.8 | 2.1 | fold enrichment | 14.2 | 11.4 | fold enrichment | 1.6 | 5.7 |

| All genes | Protein coding | lncRNAs | UP DEGs | Protein coding | lncRNAs | DN DEGs | Protein coding | lncRNAs |
|---|---|---|---|---|---|---|---|---|
| Chr9 | 181.0 | 250.5 | Chr9 | 28054.7 | 35068.3 | Chr9 | 2377.5 | 46757.8 |
| Cluster 128-138 | 48.5 | 113.6 | Cluster 128-138 | 2000.0 | 3333.3 | Cluster 128-138 | 1428.6 | 10000.0 |
| fold enrichment | 3.7 | 2.2 | fold enrichment | 14.0 | 10.5 | fold enrichment | 1.7 | 4.7 |

| All genes | Protein coding | lncRNAs | UP DEGs | Protein coding | lncRNAs | DN DEGs | Protein coding | lncRNAs |
|---|---|---|---|---|---|---|---|---|
| Chr10 | 186.0 | 194.5 | Chr10 | 27074.9 | 33843.7 | Chr10 | 2115.2 | 67687.4 |
| Cluster 70-80 | 178.6 | 142.9 | Cluster 70-80 | 5000.0 | 3333.3 | Cluster 70-80 | 2000.0 | - |
| fold enrichment | 1.0 | 1.4 | fold enrichment | 5.4 | 10.2 | fold enrichment | 1.1 | - |

| All genes | Protein coding | lncRNAs | UP DEGs | Protein coding | lncRNAs | DN DEGs | Protein coding | lncRNAs |
|---|---|---|---|---|---|---|---|---|
| Chr11 | 102.7 | 168.5 | Chr11 | 13445.2 | 14939.2 | Chr11 | 1948.6 | 67226.2 |
| Cluster 62-72 | 38.8 | 85.5 | Cluster 60-70 | 1666.7 | 2500.0 | Cluster 60-70 | 909.1 | 10000.0 |
| fold enrichment | 2.7 | 2.0 | fold enrichment | 8.1 | 6.0 | fold enrichment | 2.1 | 6.7 |

| All genes | Protein coding | lncRNAs | UP DEGs | Protein coding | lncRNAs | DN DEGs | Protein coding | lncRNAs |
|---|---|---|---|---|---|---|---|---|
| Chr16 | 104.1 | 104.7 | Chr16 | 3701.1 | 14804.5 | Chr16 | 2612.6 | 88827.3 |
| Cluster 0-10 | 147.1 | 232.6 | Cluster 0-10 | 909.1 | 3333.3 | Cluster 0-10 | 2000.0 | 10000.0 |
| fold enrichment | 0.7 | 0.5 | fold enrichment | 4.1 | 4.4 | fold enrichment | 1.3 | 8.9 |
